# The O_2_-independent pathway of ubiquinone biosynthesis is essential for denitrification in *Pseudomonas aeruginosa*

**DOI:** 10.1101/2020.03.22.002311

**Authors:** Chau-Duy-Tam Vo, Julie Michaud, Sylvie Elsen, Bruno Faivre, Emmanuelle Bouveret, Frédéric Barras, Marc Fontecave, Fabien Pierrel, Murielle Lombard, Ludovic Pelosi

**Author notes:** C.-D.-T.V. and J.M. contributed equally to the work.

## Abstract

Many proteobacteria, such as *Escherichia coli*, contain two main types of quinones, benzoquinones represented by ubiquinone (UQ) and naphthoquinones such as menaquinone (MK) and dimethyl-menaquinone (DMK). MK and DMK function predominantly in anaerobic respiratory chains, whereas UQ is the major electron carrier used for reduction of dioxygen. However, this division of labor is probably not so stric. Indeed, a pathway that produces UQ under anaerobic conditions in an UbiU-, UbiV- and UbiT-dependent manner has been recently discovered in *E. coli* while its physiological relevance is not yet understood because of the presence of MK and DMK in this bacterium. In the present study, we established that UQ_9_ is the single quinone of *P. aeruginosa* and that is required for growth under anaerobic respiration (denitrification). We demonstrated that ORFs *PA3911, PA3912* and *PA3913*, which are homologues to the *E. coli ubiT, ubiV* and *ubiU* genes, respectively, were essential for UQ_9_ biosynthesis and thus for denitrification in *P. aeruginosa*. These three genes were hereafter called *ubiT_Pa_, ubiV_Pa_* and *ubiU_Pa_*. We showed that UbiV_Pa_ accommodates a [4Fe-4S] cluster. Moreover, we demonstrated that UbiU_Pa_ and UbiT_Pa_ were able to bind UQ and that the isoprenoid tail of UQ was the structural determinant for the recognition by these Ubi proteins. Since the denitrification metabolism of *P. aeruginosa* is believed to be important for pathogenicity in cystic fibrosis patients, our results highlight the O_2_-independent UQ biosynthesis pathway as a new possible target to develop innovative antibiotics.

## INTRODUCTION

The opportunistic pathogen *Pseudomonas aeruginosa* has a remarkable ability to grow under a variety of environmental conditions such as soil and water as well as animal-, human-, and plant-host-associated environments. *P. aeruginosa* is responsible for numerous acute and chronic infections and poses a major health risk for patients with severe burns, cystic fibrosis (CF) or in a severely immunocompromised states (1,2).

The utilization of various carbon sources and energy metabolism (respiration or fermentation) might contributes to the environmental adaptation of *P. aeruginosa* (3). Its main energy-producing system is respiration, which requires a proton motive force (pmf) used for ATP synthesis. The pmf is produced by the transfer of electrons and protons from reduced donors to oxidized acceptors *via* the quinone pool. Whereas the dehydrogenases and reductases involved in respiratory metabolism have been well described and annotated in the genome of *P. aeruginosa* PAO1 (4,5), the composition of its quinone pool has not yet fully established yet in this bacterium. Studies in the sixties suggested ubiquinone 9 (UQ_9_) as a major quinone of aerobically-grown *P. aeruginosa* (6) and UQ_9_ is therefore believed to be essential for aerobic respiration (7).

Proteobacteria contain two main types of quinones, benzoquinones and naphthoquinones, represented respectively by UQ (or coenzyme Q) / plastoquinone (PQ) and menaquinone (MK) / demethylmenaquinone (DMK) (8). Typically, MK and DMK function predominantly in anaerobic respiratory chains, whereas UQ and PQ are the major electron carriers used for reduction of dioxygen by various cytochrome oxidases (8). Recent data indicated that the metabolic use of various quinone species according to environmental dioxygen availability might be more complex than initially thought. Indeed, using *E. coli* as a model, we highlighted a pathway conserved across many bacterial species and able to produce UQ under anaerobic conditions (9). The classical UQ biosynthesis pathway requires O_2_ for three hydroxylation steps (10). Obviously, the flavin-dependent monooxygenases UbiI, UbiF, and UbiH, which catalyze the O_2_-dependent hydroxylation steps are not involved in the anaerobic pathway, nor are the accessory UbiK and UbiJ proteins implicated in the assembly and/or stability of the aerobic Ubi-complex (11). Seven enzymes (UbiA, B, C, D, E, G and X) catalyzing the prenylation, decarboxylation and methylation of the phenyl ring of the 4-hydroxybenzoate precursor are common to both pathways (9). In addition, the anaerobic pathway requires UbiT, UbiU and UbiV proteins (9). However, as explained above, the metabolic relevance of the O_2_-independent UQ pathway is not yet clearly understood to date.

In absence of O_2_, *P. aeruginosa* is able to carry out anaerobic respiration with nitrate and nitrite as terminal electron acceptors of the respiratory chain. This process called denitrification allows the reduction of soluble nitrate (NO_3_^-^) and nitrite (NO_2_^-^) to gaseous nitrous oxide (N_2_O) or molecular nitrogen (N_2_) (12). Because *P. aeruginosa*-infected mucus in CF airway is depleted of oxygen and enriched in nitrate and nitrite, the anaerobic metabolism of *P. aeruginosa* via the denitrification pathway is believed to be important for its pathogenicity (13). Four sequential reactions involving metalloenzymes are needed to reduce nitrate to N_2_, i.e. nitrate reductase, nitrite reductase, nitric oxide reductase and nitrous oxide reductase. *P. aeruginosa* was considered as a paradigm of the denitrification pathway and all the reductases involved in this metabolism have been widely studied as well as the regulatory network controlling the denitrification genes (3,4,14). However, the anaerobic quinone pool of *P. aeruginosa* has not been characterized so far.

In the present study, we discovered that UQ_9_ is essential for the growth of *P. aeruginosa* PAO1 strain in denitrification medium. We identified in this bacterium the ORFs *PA3911, PA3912* and *PA3913* as homologs to *E. coli ubiT, ubiV* and *ubiU*, respectively. Our results showed that these three genes hereafter called *ubiT_Pa_, ubiV_Pa_* and *ubiU_Pa_* are essential components of the O_2_-independent UQ_9_ biosynthetic pathway of *P. aeruginosa*. We demonstrated that i), UbiV_Pa_ binds a [4Fe-4S] cluster and ii) UbiU_Pa_ and UbiT_Pa_ copurify with UQ by recognizing the isoprenoid tail. Such as molecular pathway for UQ production was found only in proteobacteria (9) where it can exert an essential role under anaerobic conditions, as demonstrated here. Taken together, our results highlight that this pathway could be an interesting lead for the development of antibiotics targeting the denitrification metabolism.

## RESULTS

### UQ_9_ is the single quinone of P. aeruginosa

The quinone content of *P. aeruginosa* PAO1, grown under ambient air or anaerobiosis (denitrification), was determined using electrochemical detection of lipid extracts separated by HPLC and compared to those obtained from *E. coli*. Whatever the conditions of growth, a single quinone species eluting at 11.5 min was present in the analyses of *P. aeruginosa* lipid extracts, UQ_10_ being used as standard (Fig. 1*A* and 1*B*). Mass spectrometry analysis showed a predominant ammonium adduct (M_+_NH_4_^+^) with *m/z* ratio of 812.7 (Fig. 1*C*), together with minor adducts such as Na^+^ (817.6) and H^+^ (795.7) (Fig. S1). These masses identify UQ_9_ (monoisotopic mass 794.6) as the quinone produced by *P. aeruginosa*. Membranes of *E. coli* contain UQ_8_ and naphthoquinones (DMK_8_ and MK_8_). The absence of detectable level of naphthoquinones in *P. aeruginosa* lipid extracts, with or without oxygen (Fig.1*A* and 1*B*), is in agreement with the absence of their biosynthetic pathways (MK or futalosine pathways) in *P. aeruginosa* genomes. It is also interesting to note that the UQ content of *E. coli* was higher under aerobiosis compared to anaerobiosis (97±4 *versus* 42±6 pmol of UQ_8_ per mg of cells), whereas we found the opposite for *P. aeruginosa* (95±4 *versus* 126±4 pmol of UQ_9_ per mg of cells). Together, our results establish that UQ_9_ is the single quinone of *P. aeruginosa* PAO1 and suggest that UQ_9_ is used under denitrification conditions.

**Figure 1:**
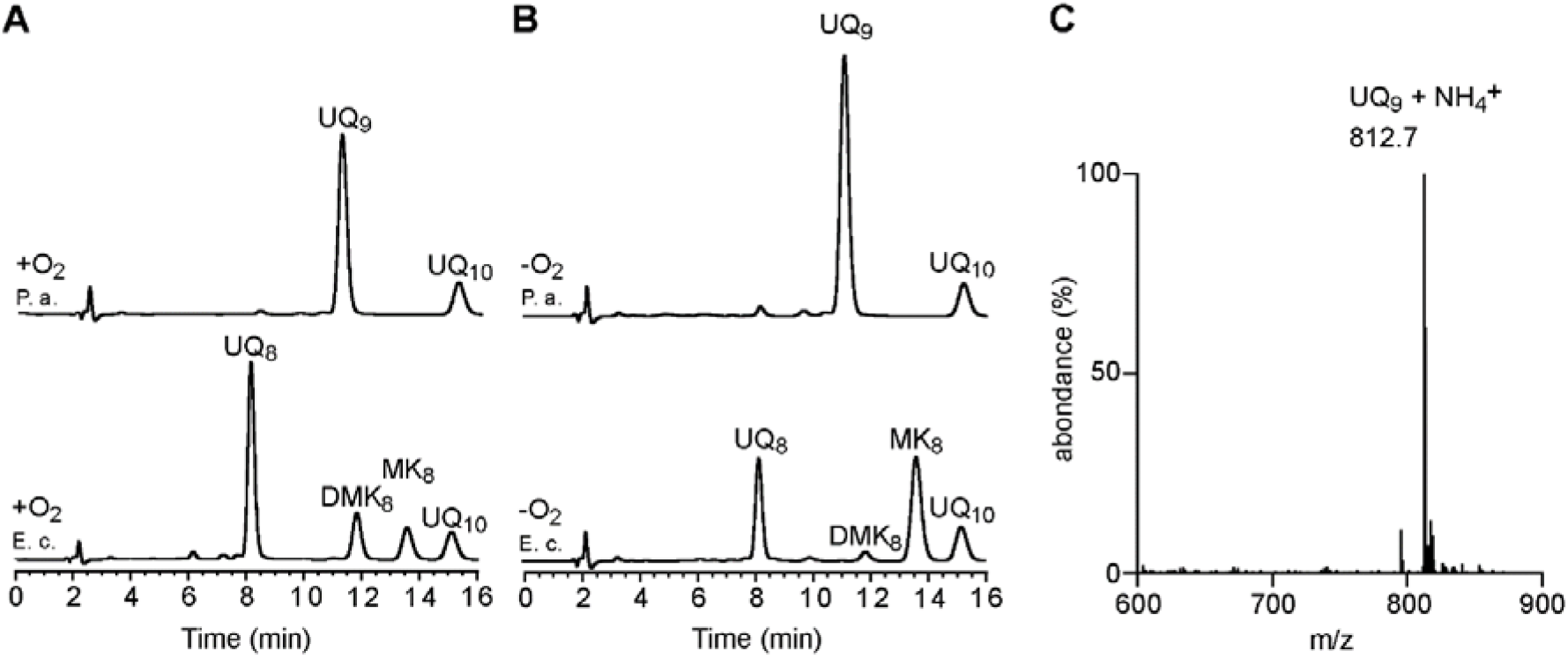
UQ_9_ is the single quinone used by *P. aeruginosa* in aerobic and anaerobic conditions. HPLC-ECD analysis of lipid extracts from 1 mg of cells after growth of *E. coli* MG1655 (E. c.) and *P. aeruginosa* PAO1 (P. a.) aerobically (+O_2_) in LB medium (*A*) or anaerobically (-O_2_) in denitrification medium (*B*). The chromatograms are representative of three independent experiments. The peaks corresponding to UQ_8_, UQ_9_, DMK_8_, MK_8_ and the UQ_10_ as a standard are indicated. *C*, Mass spectrum of the quinone eluting at 11.5 min from extracts of *P. aeruginosa* grown in aerobic and anaerobic cultures.

### Identification of ubi genes in the genome of P. aeruginosa PAO1

To identify the Ubi proteins of *P. aeruginosa*, IspB, UbiX, and UbiA to UbiK from *E. coli* MG1655 were first screened for homologs in the *P. aeruginosa* PAO1 protein sequence data set available at http://www.pseudomonas.com/ using the BLASTP software. As listed in Table S1, the analysis disclosed the presence of 11 homologous proteins (IspB, UbiA to UbiE, UbiG to UbiJ and UbiX). As reported previously, the functional homolog of UbiF is a Coq7-like hydroxylase (15) and the corresponding PA0655 protein was shown to be essential for aerobic UQ_9_ biosynthesis (16). Overall, we propose that the O_2_-dependent UQ biosynthetic pathways in *P. aeruginosa* and *E. coli* share a similar pattern (Fig. S2).

Under anaerobiosis, *E. coli* still synthesizes UQ and we recently identified three genes that we called *ubiT, ubiV* and *ubiU*, as essential for this process (9). Homologues of *ubiT, ubiV* and *ubiU* were also identified in *P. aeruginosa* PAO1 and correspond to ORFs *PA3911, PA3912* and *PA3913*, respectively (Table S1 and Fig. S2). These genes were called hereafter *ubiT_Pa_, ubiV_Pa_* and *ubiU_Pa_*. The three genes are predicted to form an operon, *ubiUVT* (www.pseudomonas.com). Interestingly, this operon is located downstream of the genes *moeA1, moaB1, moaE, moaD* and *moaC* involved in the biosynthesis of the molybdopterin cofactor (MoCo) (Fig. S3), which is essential for nitrate reductase activity (17). Next, we evaluated the metabolic relevance of the O_2_-independent UQ biosynthetic pathway in *P. aeruginosa* by studying mutants of *ubiT_Pa_, ubiU_Pa_* and *ubiV_Pa_* genes.

### Tn mutants of ubiV_Pa_ and ubiU_Pa_ present a growth defect for denitrification and an impaired UQ_9_ content

The physiological importance of the proteins UbiT_Pa_, UbiU_Pa_ and UbiV_Pa_ was first investigated using transposon (Tn) mutants PW7609 (*ubiT_Pa_*), PW7610 (*ubiV_Pa_*), PW7611 (*ubiV_Pa_*), PW7612 (*ubiU_Pa_*) and PW7613 (*ubiU_Pa_*) and the isogenic parental strain MPAO1 (wild-type strain from the Manoil collection) as control (Table S2). Aerobic growth in LB medium was similar between the Tn mutants and the wild-type strain MPAO1 (Fig. 2*A*) and the Tn mutants presented a UQ_9_ level comparable to the wild-type (Fig. 2*B*). Thus, UbiT_Pa_, UbiU_Pa_ and UbiV_Pa_ are not involved in the O_2_-dependent UQ biosynthetic pathway of *P. aeruginosa*. In contrast, the growth of the *ubiV_Pa_* and *ubiU_Pa_* mutants was severely impaired in denitrification conditions (Fig. 2*C*) and their UQ_9_ content was strongly lowered (Fig. 2*B*). These results suggest the overall requirement of *ubiU_Pa_* and *ubiV_Pa_* for denitrification in *P. aeruginosa*, supposedly *via* their involvement in O_2_-independent UQ biosynthesis. Surprisingly, the growth of the *ubiT_Pa_* Tn mutant PW7609 was not affected (Fig. 2*C*) and it showed around 40% UQ_9_ compared to the wild-type in anaerobic cultures (Fig. 2*B*). In this mutant, the Tn is inserted at the fifth base of the *ubiT_Pa_* gene, potentially leading to only partial inactivation of the gene (Table S2). We note that previous studies with *E. coli ubi* mutants showed that only 20% UQ was sufficient to maintain a wild-type growth phenotype (18,19).

**Figure 2:**
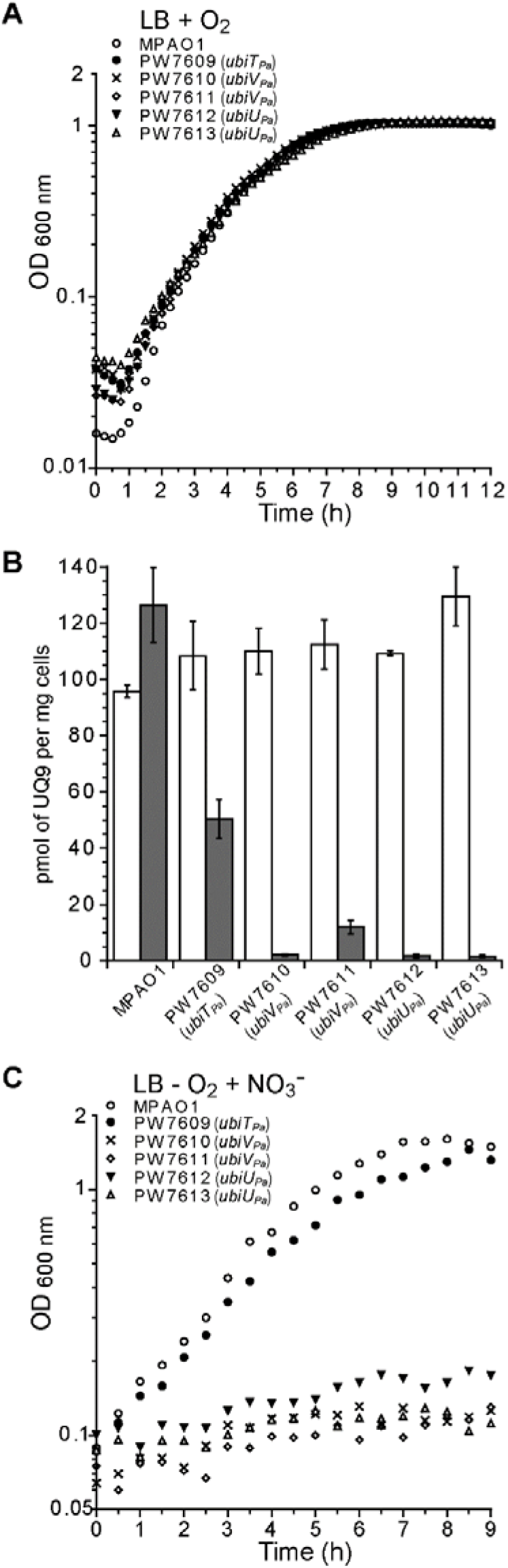
UQ_9_ content is impaired in the Tn mutants of *ubiU_Pa_, ubiV_Pa_* and *ubiT_Pa_* in denitrification medium. Representative growth curves of wild-type MPAO1 and Tn mutants grown aerobically in LB medium (*A*) or anaerobically in denitrification medium (*C*). Data are representative of three independent experiments. *B*, Quantification of cellular UQ_9_ content (*n=*3) in lipid extracts from wildtype MPAO1 and Tn mutants cells grown aerobically in LB medium (white bars) or anaerobically in denitrification medium (grey bars). Error bars represent S.D.

### Denitrification is dependent of ubiT_Pa_, ubiU_Pa_ and ubiV_Pa_ genes via their involvement in O_2_-independent UQ biosynthesis

As *ubiT_Pa_, ubiV_Pa_* and *ubiU_Pa_* are localized next to each other in the genome of PAO1, the transposon inserted in the mutants previously studied might impact the expression of the neighbouring genes. In addition, it is likely that the Tn mutant PW7609 is not disrupting properly the *ubiT_Pa_* gene. Thus, for each of the three genes, we constructed knockouts (KO) as well as complementation mutants in the parental strain PAO1. All deletion mutant strains (called hereafter *ubiTUV*-KO) shared a growth defect under denitrification coupled to a strong decrease of UQ_9_ content compared to the wild-type (Fig. 3*A* and 3B), whereas UQ_9_ content and growth were normal in aerobic conditions (Fig. 3*B* and 3*C*). Interestingly, upon complementation, bacterial growth and UQ_9_ levels were restored to those of the PAO1 strain used as control (Fig. 4*A*).

**Figure 3:**
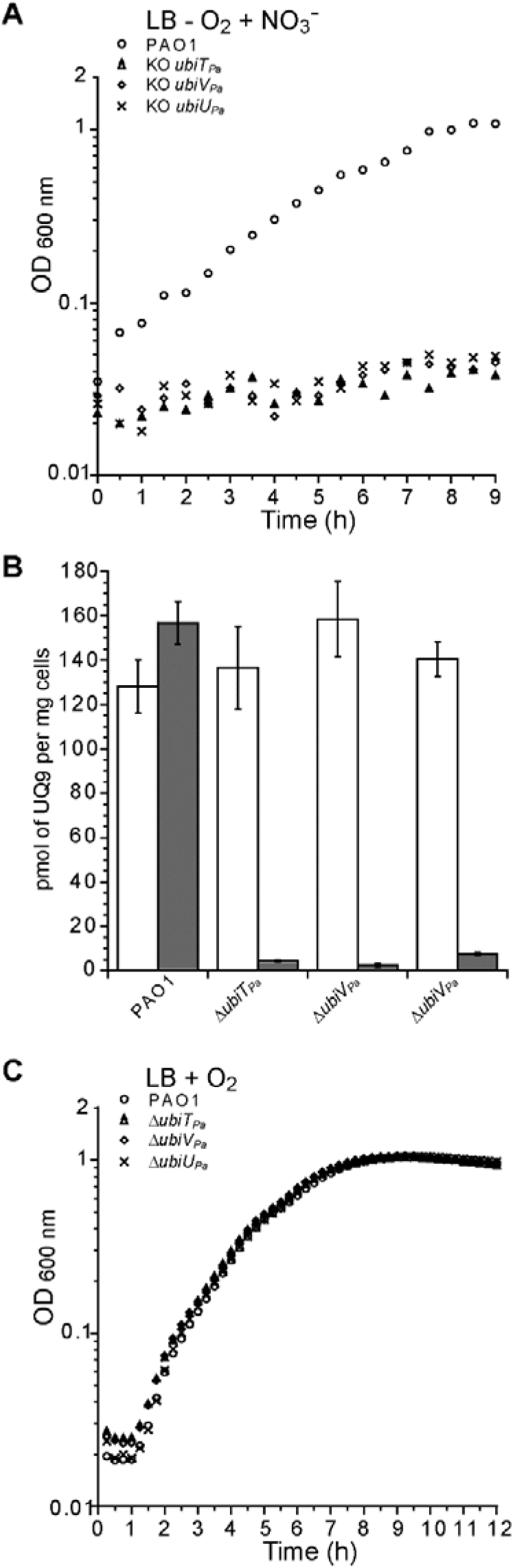
*ubiU_Pa_, ubiV_Pa_ and ubiT_Pa_* are essential genes for anaerobic UQ_9_ biosynthesis and for denitrification. Representative growth curves of wild-type PAO1 and *ubiTUV*-KO strains grown (*A*) in denitrification medium or (*C*) aerobically in LB medium. *B*, Quantification of cellular UQ_9_ content (*n=*3) in lipid extracts from wild-type PAO1 and KO cells grown aerobically in LB medium (white bars) or in denitrification medium (grey bars). Error bars represent S.D.

To confirm that UQ was directly involved in the restoration of the anaerobic growth of the *ubiTUV*-KO strains, UQ_4_ solubilized in methanol was added to the denitrification medium at 5 and 50 μM final concentrations. After 24h of anaerobic incubation, colony-forming units per mL (CFU/mL) of each KO strain were estimated and compared to the same strain cultivated without UQ_4_. The most significant results were obtained with 50 μM of UQ_4_, which increased CFU of *ubiTUV*-KO strains by 8-15 fold (Fig. 4*B*). We noted a substantial toxicity of methanol on the wild-type strain (Fig. 4*B*, Ct lane), suggesting that the positive effect of UQ_4_ on the *ubiTUV*-KO strains is likely underestimated. Taken together, our results show unequivocally that UbiT_Pa_, UbiU_Pa_ or UbiV_Pa_ are needed for denitrification *via* their involvement in UQ biosynthesis.

**Figure 4:**
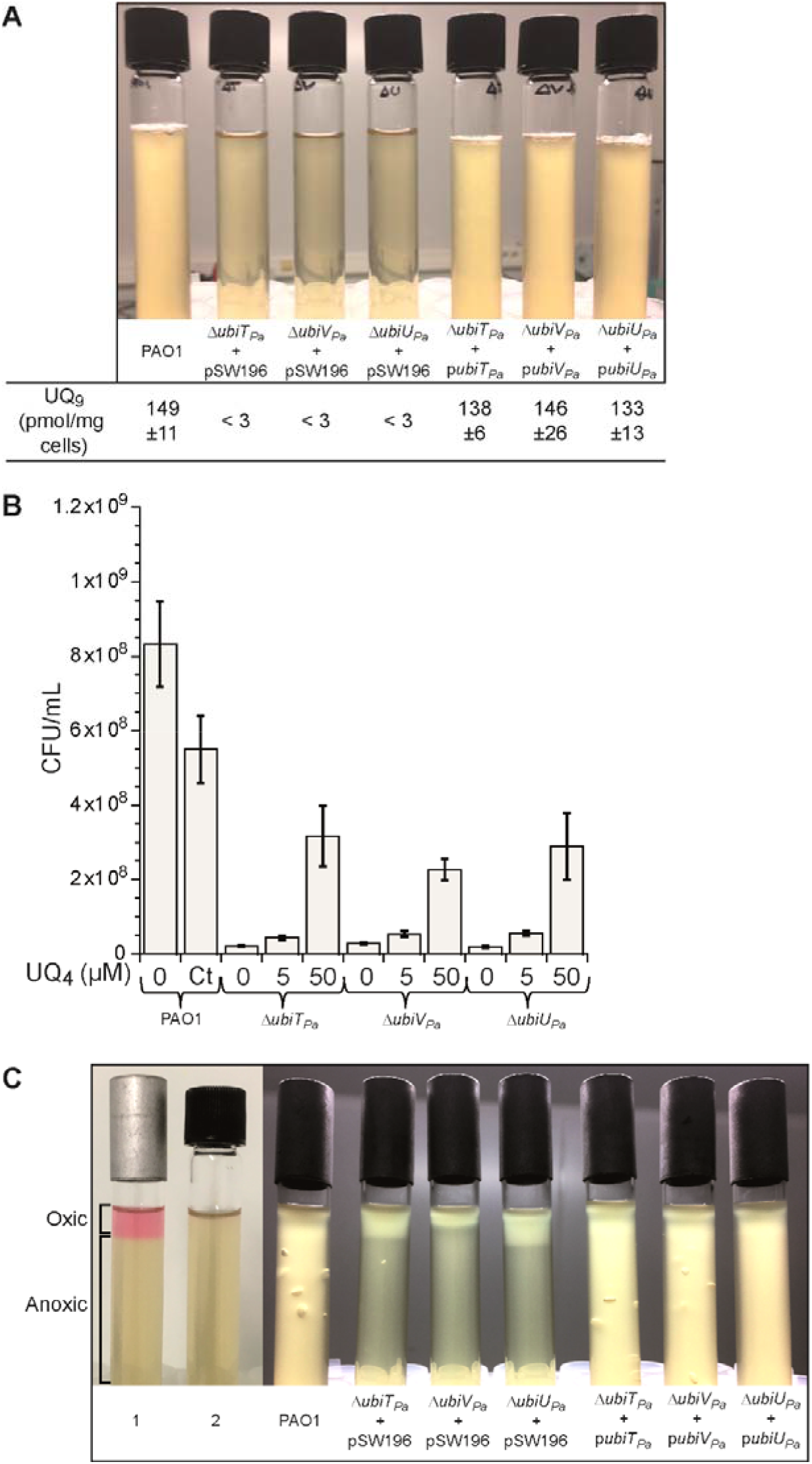
Complementation of *ubiTUV*-KO strains restores bacterial growth over the entire O_2_ range in a UQ-dependent manner. *A*, Photographs of culture tubes after overnight growth under anaerobic conditions in denitrification medium of *ubiTUV*-KO strains transformed with the empty vector pSW196 or the same vector carrying the corresponding wild-type allele (*ubiT_Pa_, ubiU_Pa_* and *ubiV_Pa_*). The parental strain PAO1 was used as a control and UQ_9_ content of wild-type and *ubiTUV*-KO strains cultured anaerobically was assayed (n=3). *B, ubiTUV*-KO strains were cultured in denitrification medium supplemented with methanol-solubilized UQ_4_ at 5 or 50 μM final concentration. After 24h of incubation, colony-forming units per mL (CFU/mL) of each KO strain were estimated and compared to the same strain grown without UQ_4_. As a control, toxicity of methanol was tested on the parental strain grown in the same medium but supplemented with 4.5% (v/v) methanol (Ct), which is the final concentration of methanol corresponding to the adding of 50 μM of UQ_4_. Data are representative of three independent experiments. *C*, As in *A*, but in soft-agar tubes after overnight culture. All the strains studied were inoculated into anaerobic tubes and then exposed to ambient air to create an oxygen gradient. The controls (Ct) correspond to soft agar tubes supplemented with 2.5 μg/mL resazurin and then incubated with (1) or without air (2). Oxic and anoxic parts of the agar are indicated. For all strains containing pSW196 vectors, denitrification medium was also supplemented with 0.1% (w/v) final concentration of arabinose to induce *P*_BAD_ promoter.

### UbiT_Pa_, UbiU_Pa_ and UbiV_Pa_ are needed for the process of denitrification

We used soft-agar experiments to examine dioxygen and nitrate requirements of *ubiTUV*- KO strains with or without the wild-type allele. Soft agar was prepared anaerobically in LB medium containing KNO_3_ at 100 mM final concentration and then exposed to ambient air. Oxygen diffuses through the agar to form a gradient, the highest concentration being at the top of the agar (Fig. 4*C*, lane 1). As shown in Fig. 4*C*, parental strain PAO1 and complemented *ubiTUV*-KO strains grew throughout the tube because they were able to use aerobic respiration as well as denitrification. In contrast, the growth of the *ubiTUV*-KO strains harboring the empty vector was restricted to the oxygenated part of the medium, whereas the presence of the respective genes on the plasmids allowed growth in the anaerobic medium (Fig. 4*C*). The bubbles observed in the soft agar correspond to gas evolution of N_2_O and/or N_2_ (12), suggesting a restoration of the denitrification process in the lower part of the tube. Taken together, these results point to the requirement for *ubiT_Pa_, ubiU_Pa_* and *ubiV_Pa_* beyond the nitrate reduction step and support that UQ is probably essential for the entire denitrification process in *P. aeruginosa*.

### Molybdopterin cofactors are not involved in anaerobic UQ_9_ biosynthesis

As mentioned previously, the *ubiUVT* operon is located downstream of the genes *moeA1, moaB1, moaE, moaD* and *moaC* involved in MoCo biosynthesis. Currently, MoCo-containing hydroxylases constitute the only family known to catalyze O_2_-independent hydroxylation reactions (20). Since three O_2_-independent hydroxylation reactions are needed to synthesize UQ anaerobically (9), we reasoned that MoCo might be involved in this process, providing a rationale for the co-localization of the *ubi* and *moa/moe* genes. To test this hypothesis, we evaluated the ability of Tn mutants PW7614, PW7615/PW7616, PW7618/PW2470, PW7619/PW1920 and PW7621/PW7622 (Table S2) corresponding respectively to Tn insertions in the ORFs *moeA1, moaB1* (two mutants), *moaE* (two mutants), *moaD* (two mutants) and *moaC* (two mutants) to synthesize UQ_9_ without O_2_. As expected, all the Tn mutants exhibited a growth defect in denitrification, since MoCo is an essential component of nitrate reductase (17). However, anaerobic growth was rescued by addition of arginine as previously described (21), and we were therefore able to measure the UQ content of the cells in these conditions (Fig. S4). The UQ_9_ content of the MoCo Tn mutants was comparable to that of the wild-type strain (Fig. S4), suggesting that MoCo is not involved in anaerobic UQ_9_ biosynthesis.

### Recombinant UbiV_Pa_ is an air-sensitive [4Fe-4S] cluster-containing protein

To gain insights into their biochemical properties, we produced and purified the three proteins in *E. coli*, UbiV_Pa_ being the most soluble. First, we showed that UbiV_Pa_ purified by size exclusion chromatography (SEC) behaved as a monomer (Fig. S5*A* and S5*B*). Moreover, we noticed that the fraction containing the purified protein was slightly pink-colored with a UV-visible absorption spectrum characteristic of the presence of iron-sulfur species (22), with a band at 410 nm and broad and low intensity shoulders between 450 and 600 nm (Fig. 5*A*, dotted line) (23). However, the amount of iron and sulfur (0.22 iron and 0.22 sulfur/monomer) was largely substoïchiometric, suggesting a degradation of the [Fe-S] cluster during the purification of the protein under aerobiosis, as already observed for many other Fe-S proteins. Consistent with this hypothesis, anaerobic reconstitution of the [Fe-S] cluster allowed to obtain a brown color protein with a UV-visible spectrum displaying one broad absorption band at 410 nm, which is characteristic of a [4Fe-4S]^2+^ cluster (Fig. 5*A*, solid line) (24). The iron and sulfide determination yielded 3.90 ± 0.03 iron and 3.40 ± 0.20 sulfur/monomer of UbiV_Pa_, consistent with the presence of one [4Fe-4S] cluster/protein (Table 1). As shown in Fig. S5*C*, the [Fe-S] cluster of UbiV_Pa_ was sensitive to air.

**Figure 5:**
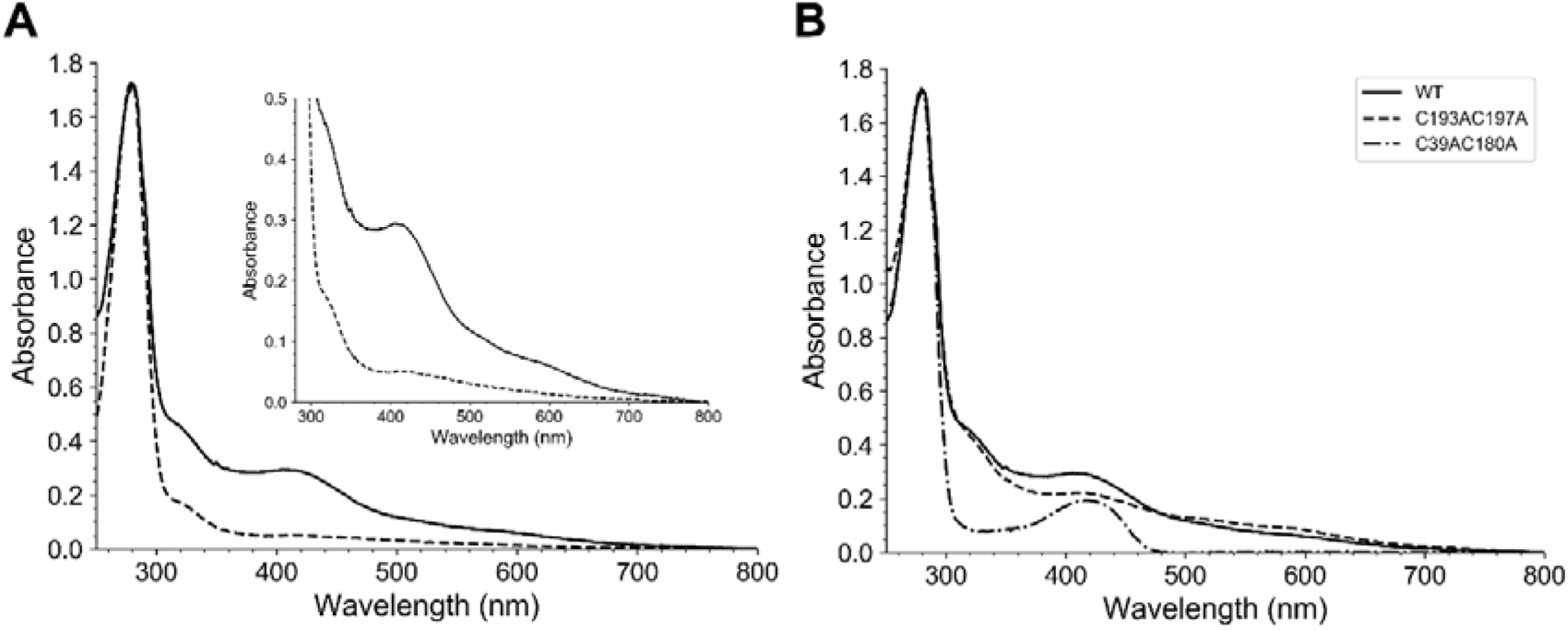
Recombinant UbiU_Pa_ is a [4Fe-4S] cluster-containing protein. *A*, UV-visible absorption of as-purified UbiU_Pa_ (dotted line, 32.6 μM) and reconstituted holo-UbiU_Pa_ (solid line, 22.7 μM). The inset is an enlargement of the 300 to 800 nm region. The molar extinction coefficient, ε_410nm_ was determined to be 12.95 ± 0.5 mM^-1^ cm^-1^ for holo-UbiV_Pa_. *B*, Comparative UV-visible absorption spectra of wild-type and different Cys-to-Ala mutants of UbiV_Pa_ after metal cluster reconstitution. Proteins were analyzed at the following concentrations: 22.7μM WT, 34.8 μM C39AC180A and 15.9 μM C193AC197A. Proteins were suspended in buffer 50 mM Tris-HCl, 25 mM NaCl, 15% (v/v) glycerol, 1 mM DTT, pH 8.5.

**Table 1:**
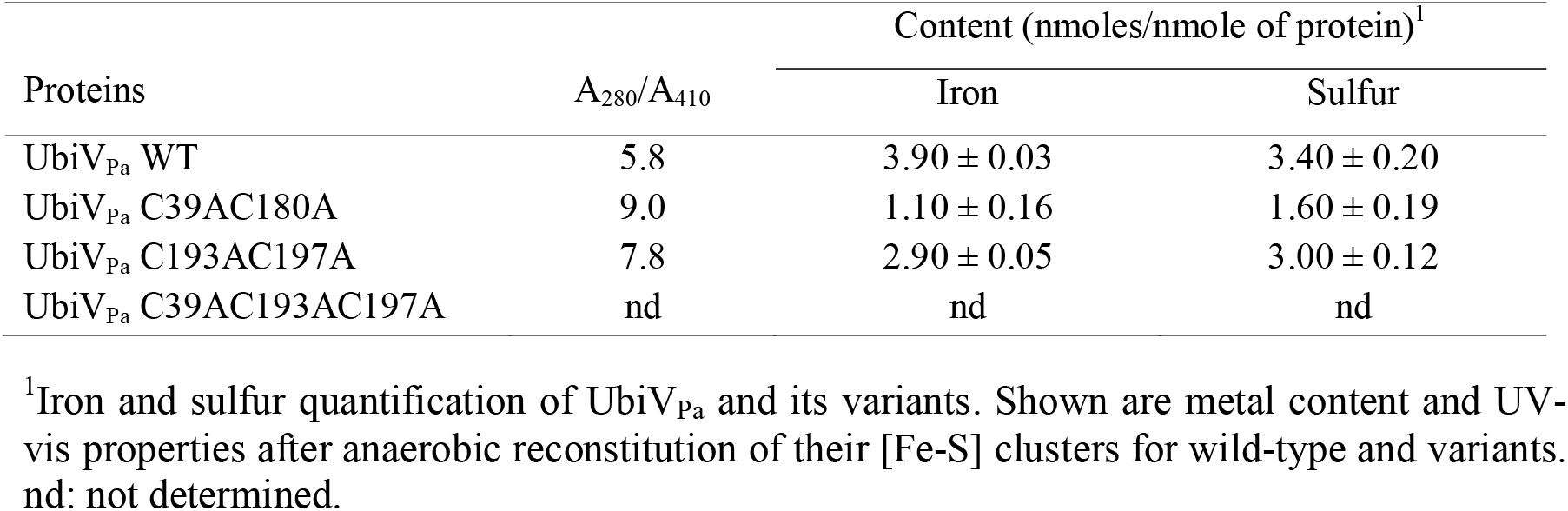
Spectral characterization of UbiV_Pa_ and its variants.

| Proteins | A<sub>280</sub>/A<sub>410</sub> | Content (nmoles/nmole of protein)<sup>1</sup> |  |
| --- | --- | --- | --- |
|  |  | Iron | Sulfur |
| UbiV<sub>Pa</sub> WT | 5.8 | 3.90 ± 0.03 | 3.40 ± 0.20 |
| UbiV<sub>Pa</sub> C39AC180A | 9.0 | 1.10 ± 0.16 | 1.60 ± 0.19 |
| UbiV<sub>Pa</sub> C193AC197A | 7.8 | 2.90 ± 0.05 | 3.00 ± 0.12 |
| UbiV<sub>Pa</sub> C39AC193AC197A | nd | nd | nd |

<sup>1</sup>Iron and sulfur quantification of UbiV<sub>Pa</sub> and its variants. Shown are metal content and UV-vis properties after anaerobic reconstitution of their [Fe-S] clusters for wild-type and variants. nd: not determined.

Four strictly conserved cysteines (C39, C180, C193, and C197) arranged in a CX_n_CX_12_CX_3_C motif (X representing any amino acid) are found in UbiV_Pa_ (9). To test if these four cysteines are important for the chelation of the [4Fe-4S] cluster present in UbiV_Pa_, we generated two double mutants (C39AC180A and C193AC197A) and a triple mutant (C39AC193AC197A). All these mutants were colorless after purification under aerobic purification and did not show any absorption band in the 350- to −550-nm region of their UV-Vis spectra (Fig. S5*D*), suggesting that they were impaired in their capacity to accommodate a [Fe-S] cluster. After reconstitution under anaerobic conditions, UbiV_Pa_ C39AC193AC197A precipitated and its UV-vis spectrum could not be recorded. Although they also had a tendency to aggregate, 10% of the double mutants behaved as monomers permitting some assays. Overall, their absorbance at 410 nm (Fig. 5*B*) and their iron and sulfur contents (Table 1) were largely decreased compared to the wild-type protein, suggesting that the four conserved cysteines are good candidates as ligands of the [4Fe-4S] cluster present in UbiV_Pa_.

### Recombinant UbiU_Pa_ and UbiT_Pa_ co-purify with UQ_8_ in E. coli

We have recently demonstrated that isoprenoid quinones were able to co-elute with the Ubi-proteins such as UbiJ (11) and that UbiT exhibits a sterol carrier protein 2 (SCP2) domain (9). To that end, we performed lipid content analysis of the UbiT_Pa_, UbiU_Pa_ and UbiV_Pa_ fractions purified from *E. coli* extracts. No isoprenoid quinones were detected co-eluting with UbiV_Pa_ (Table S3). In contrast, UQ_8_ and DMQ_8_ (2-octaprenyl-3-methyl-6-methoxy-1,4-benzoquinone) were shown to co-purify with UbiU_Pa_. This protein was purified only in the presence of detergent, as it was insoluble without it. After a two-step purification protocol, including a Ni-NTA chromatography and a SEC, the solubilized protein had still a tendency to form different oligomeric states covering the fractions 14 to 44 as shown in Fig. 6*A* and 6*B*. UQ_8_ and DMQ_8_ were mainly detected in the elution fractions 33 to 45 (Fig. 6*A*) corresponding to only a proportion of the purified UbiU_Pa_ (Fig. 6*B*). The highest contents, i.e. 488.17 pmoles of UQ_8_ per mg of protein and 19 932 AU of DMQ_8_ per mg of protein, were assayed in the fraction 39 and 40, respectively (Table S3). This corresponds to a UQ_8_/protein ratio of 1.5%. Taken together, these results show that UQ_8_ and DMQ_8_ co-purify with UbiU_Pa_ depending of its oligomerization state.

**Figure 6:**
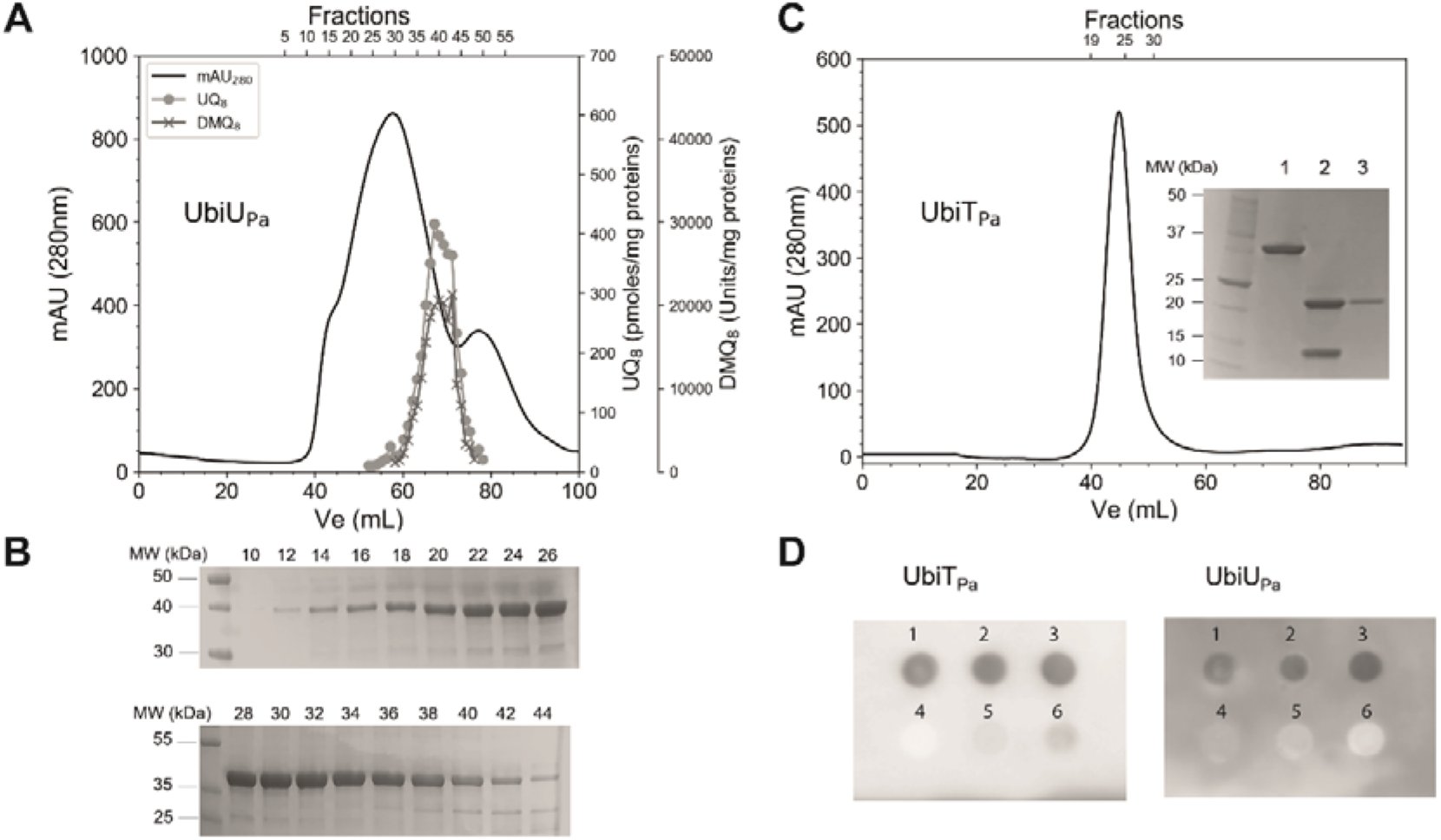
Recombinant UbiU_Pa_ and UbiT_Pa_ bind UQ_8_. *A*, Elution profile of UbiU_Pa_, 70 mg of protein were loaded on a Superdex 200 16/60 chromatography column. A quantification of UQ_8_ and DMQ_8_ in each fraction was performed by HPLC-ECD MS. A recovery of 73 and 76%, respectively for UQ_8_ and DMQ_8_, was calculated from the total content of all fractions compared with content of the UbiU_Pa_ purified fraction deposited on the Superdex 200 column. *B*, Fractions 10 to 44 analyzed by SDS-PAGE for purity. *C*, Elution profile of UbiT_Pa_ on a Superdex 200 16/60 column. Inset, SDS-PAGE, lane 1, 32 kDa TrxA-UbiT_Pa_ fusion protein, lane 2, after digestion with thrombin (UbiT_Pa_, 19.6kDa and TrxA, 12.1 kDa), lane 3, pooled fractions 20-30 of UbiT_Pa_. Quantification of UQ_8_ (pool of fractions 20 to 30) was performed by HPLC-ECD MS. *D*, Protein-lipid overlay assay between UbiU_Pa_ and UbiT_Pa_ and different lipid ligands. 2 μL of six different lipid/compound potential candidates (1, UQ_8_; 2, UQ_10_; 3, solanesol; 4, 3-methylcatechol; 5, cholesterol; 6, POPE) at 20 mM final concentration were spotted on a PVDF membrane and then incubated with UbiT_Pa_ or UbiU_Pa_ (both proteins at 0.2 μg/mL final concentration). Detection of bound proteins was performed by chemiluminescence, as described in Experimental Procedures section. Mw: molecular weights.

Due to a lack of solubility, UbiT_Pa_ was overproduced with *E. coli* thioredoxin (TrxA) as a gene fusion partner, as previously described (25). After the first step of the purification process (Ni-NTA chromatography), the 32 kDa TrxA-UbiT_Pa_ fusion protein was digested with thrombin. Then in order to remove the TrxA His-tagged protein, UbiT_Pa_ was purified by Ni-NTA chromatography coupled to SEC, as a high oligomeric form (Fig. 6*C*). The purified UbiT_Pa_ (pool of fractions 20 to 30) contained 9.75±4.57 pmoles of UQ_8_ per mg of protein (Table S3), which corresponds to a UQ_8_/protein ratio of about 0.03%.

### Recombinant UbiU_Pa_ and UbiT_Pa_ bind the isoprenoid tail of UQ

To further confirm the ability of UbiT_Pa_ and UbiU_Pa_ to bind UQ, a protein–lipid overlay assay was performed (Fig. 6*D*). We checked the possibility of these proteins to recognize UQ_10_, UQ_8_, solanesol, 3-methylcatechol, 1-palmitoyl-2-oleoyl-*sn*-glycero-3-phosphoethanolamine (POPE) and cholesterol. Solanesol is a non-cyclic terpene alcohol that consists of nine isoprene units as found in UQ_9_. 3-methylcatechol was chosen to mimic the head group of UQ. POPE is the major lipid component of the inner membrane of *E. coli* (26). Finally, cholesterol was used as a sterol standard. Fig. 6*D* shows that UbiT_Pa_, and UbiU_Pa_ did not interact with 3-methylcatechol, POPE, or cholesterol in our experimental conditions. In contrast, both proteins were able to recognize UQ_10_, UQ_8_ and solanesol. We established the ability of UbiT_Pa_ to bind phosphatidic acid (PA) as previously demonstrated by Groenewold *et al*. (25) (Fig. S6). Together, we show that UbiT_Pa_ and UbiU_Pa_ are able to bind the isoprenoid tail of UQ in agreement with their involvement in the O_2_-independent biosynthesis pathway of UQ.

## DISCUSSION

UQ acts as a membrane-embedded electron and proton shuttle and is a key molecule in the respiratory metabolism of proteobacteria. Biosynthesis of UQ in aerobic conditions has been widely studied and includes a series of enzymatic reactions in which a benzene ring undergoes a series of modifications involving a prenylation, a decarboxylation, three methylations, and three hydroxylations (27). Our chemical analysis identified UQ_9_ as the single quinone in the membranes of aerobically grown *P. aeruginosa* cells, which is in agreement with the literature (7,16). We found in *P. aeruginosa* homologues of the genes known to be involved in UQ biosynthesis in *E. coli*, except *ubiF* (Table S1). Indeed, as already published, *P. aeruginosa* exhibits a yeast COQ7 protein homolog, which catalyzes the same reaction as UbiF from *E. coli* (15,16). Thereby, both bacteria share a similar UQ biosynthesis pathway involving three hydroxylases - UbiI, UbiH and UbiF or COQ7 - using O_2_ as a co-substrate (Fig. S2). As no other isoprenoid quinone was detected in lipid extracts of *P. aeruginosa*, we suppose that UQ_9_ is essential for aerobic growth of this bacterium.

In the absence of oxygen, *P. aeruginosa* can grow by dissimilatory nitrate respiration by using nitrate or nitrite as alternative terminal electron acceptors of the respiratory chain. This metabolic process, known as denitrification, has been widely studied (3). However, the component acting to transfer electrons from primary dehydrogenases to nitrate or nitrite reductases was not clearly identified to date (3). In the present study, we identified UQ_9_ as the single quinone synthesized in anaerobic conditions, suggesting that *P. aeruginosa* possesses an O_2_-independent UQ biosynthesis pathway. We recently identified such pathway in *E. coli* (9). Here, we characterized three genes, *ubiT_Pa_ (PA3911), ubiU_Pa_ (PA3913)* and *ubiV_Pa_ (PA3912) as* essential for the anaerobic UQ_9_ biosynthesis in *P. aeruginosa* and dispensable for the aerobic one. These genes are homologs to the *ubiT, ubiU* and *UbiV* previously identified in *E. coli*, which grows normally in anaerobiosis in a UQ-independent manner because of the presence of naphthoquinones (9) (28) (29). In contrast, we demonstrated that these three genes were essential to anaerobic denitrification metabolism of *P. aeruginosa*, which is in agreement with the presence of a single quinone corresponding to UQ. In line with our results, *ubiV_Pa_* (*PA3912*) and *ubiU_Pa_* (*PA3913*) are expressed in response to anaerobic conditions (30) and the abundance of UbiU_Pa_ protein was increased during anaerobic growth (31). Moreover, using random transposon mutagenesis, *ubiV_Pa_* (*PA3912*) and *ubiU_Pa_* (*PA3913*) were already reported as essential for anaerobic growth of *P. aeruginosa* on nitrate and nitrite as alternative terminal electron acceptors of the respiratory chain (21).

Although its contribution is still poorly understood for within-host growth, anaerobic respiration of *P. aeruginosa* is likely to be significant for promoting virulence mechanisms in chronic lung infections (13). Indeed, according to O_2_ assays directly in the lungs of CF patients, the infected endobronchial mucus is subject to severe hypoxia or even anoxia (32). Two hypotheses were proposed to explain this phenomenon in the literature. First, the biofilm structure *per se* may establish an oxygen gradient across. However, this hypothesis is currently questioned due to the strong morphological differences between *in vivo* and *in vitro* biofilm production, the latter being much more thicker than the former, for example (13). The second hypothesis is that accelerated O_2_ consumption in the biofilm may result from activated polymorphonuclear leukocytes that produce superoxide (33) and to a lesser extend nitric oxide (34). Additional host responses also affect the availability of alternative electron acceptors for anaerobic bacterial metabolism. Indeed, high levels of nitrate and nitrite have been measured in sputum from CF patients (35). From all these observations and as UQ is an essential component of the denitrification metabolism in *P. aeruginosa*, we propose that UbiT_Pa_, UbiV_Pa_ and UbiU_Pa_ may contribute to the CF lung infection in patients (work in progress in our laboratory). This hypothesis is supported by a recent quantitative proteomics approach revealing increased abundance of the three proteins in anaerobic biofilms grown under conditions of the cystic fibrosis lung (25). Moreover, deduced from a high-throughput sequencing of Tn libraries from *P. aeruginosa* strain PA14, it appears that homologs of *ubiA, ubiB, ubiC, ubiE* or *ubiT* genes were found to be essential for this bacterium to colonize the murine gastrointestinal tract (36). O_2_-dependent and O_2_-independent UQ biosynthesis pathways share the proteins UbiA, UbiB, UbiC and UbiE (9). However, homologs of UbiT are only required under anaerobic conditions (9 and this work), which suggests that O_2_-independent UQ biosynthesis could be essential for bacterial virulence. This hypothesis is supported by the essential contribution of UbiU and UbiV homologs to *Yersinia ruckeri* virulence (37).

In order to better understand their role, we decided to overproduce in *E. coli*, purify and biochemically characterize UbiT_Pa_, UbiU_Pa_ and UbiV_Pa_. Our results showed that recombinant UbiV_Pa_ is an air-sensitive Fe-S containing protein, as UbiV from *E. coli* (9), and we demonstrated that cysteines 39, 180, 193 and 197 were ligands to the [4Fe-4S] cluster found in UbiV_Pa_. These results confirm the conservation of a four cysteines pattern coordinating a Fe-S cluster across homologs of UbiV. This patter is also found in RlhA and TrhP (38,39), two proteins that belong to the same proteases U32 family than UbiU and UbiV, and that are also involved in O_2_-independent hydroxylation reactions in *E. coli* (38,39). However, the function of the iron-sulfur centers in the hydroxylation mechanism remains to be understood.

As a member of the U32 protease family, UbiU_Pa_ also presents four conserved cysteines (C169, C176, C193 and C232). Unfortunately, we failled to reconstitute a Fe-S cluster without success and obtained instead protein precipitation. Indeed, UbiU_Pa_ is an unstable protein. Unlike UbiU from *E. coli*, which forms a stable heterodimer UbiU-UbiV complex (9), we were not able to solubilize UbiU_Pa_ by coproducing it with its potential partner UbiV_Pa_. The fact that we produced *P. aeruginosa* proteins in *E.coli* could explain this difference in behavior. Nevertheless, an UbiU-UbiV complex in *P. aeruginosa* remains reasonable hypothesis that needs further investigations. We were able to purify UbiU_Pa_ alone with significant quantities of UQ_8_ and DMQ_8_, whereas purified UbiV_Pa_ contained no quinones. This result was confirmed by a protein-lipid overlay assay, which showed that the isoprenoid tail of UQ was the structural determinant for the recognition by UbiU_Pa_. From these results, we propose that UbiU would bind UQ and reaction intermediates of the anaerobic UQ pathway.

Homologs of UbiT and UbiJ contain a SCP2 domain (9,40), involved in protein-lipid interactions and UbiJ from *E. coli* co-purified with UQ_8_ (11). Moreover, a previous study showed that PA3911 (UbiT_Pa_) was able to bind specifically PA, the central hub of phospholipid metabolism (25). Here we showed that UbiT_Pa_ binds to UQ_8_ and shares with UbiU_Pa_ the recognition of the isoprenoid tail of UQ. Taken together, these results support the hypothesis that UbiT may be the counterpart of UbiJ in anaerobic conditions. We propose that UbiT would bind UQ intermediates and would stabilize a putative anaerobic Ubi complex that has yet to be demonstrated.

## EXPERIMENTAL PROCEDURES

### Bacterial strains and growth conditions

*P. aeruginosa* and *E. coli* strains used in this study are listed in Table S2. We obtained the collection of transposon (Tn) mutants in *P. aeruginosa* MPAO1 strain from the Manoil Laboratory, Department of Genome Science, University of Washington, USA (41,42). The Tn insertion site of the mutant strains were verified by sequencing (GATC Biotech, Konstanz, Germany) using PCR primers recommended by the library curators. *P. aeruginosa* strains were aerobically maintained at 37°C on lysogeny broth (LB) agar plates. For quinone assay, aerobic cultures (5 mL) were performed in LB medium at 37°C with rotary shaking at 200 rpm. Anaerobic growth of *P. aeruginosa* were performed in a 12 mL Hungate tubes containing LB medium supplemented with KNO_3_ as an electron acceptor (100 mM final concentration) (43), hereafter called denitrification medium, and deoxygenated 30 min by argon bubbling (O_2_<0.1 ppm) prior to autoclaving. In some experiments, LB medium was supplemented with arginine at a final concentration of 40 mM instead of KNO_3_ (44). Hungate tubes were inoculated through the septum with 100 μL of overnight cultures taken with disposable syringe and needles from closed Eppendorf tubes filled to the top. Cultures in hungate tubes were used for measuring the quinone contents. For aerobic growth studies, aerobic overnight cultures were used to inoculate a 96-well plate to obtain a starting optical density at 600 nm (OD_600_) of 0.05 and further incubated with shaking at 37°C. Changes in OD_600_ were monitored every 10 min for 12 h using the Infinite 200 PRO microplate reader (Tecan, Lyon, France). For anaerobic growth curve studies, overnight cultures in 50 mL closed tubes of not degassed denitrification medium were used to inoculate 400 mL bottles to obtain a starting OD_600_ of 0.05. Then, bacteria were grown anaerobically by sparging argon (O_2_<0.1 ppm) and bacterial cultures were monitored spectrophotometrically (OD_600_) at 30 min intervals for 9h. *E. coli* MG1655 and DH5α were grown on LB agar or in LB liquid. When required, the medium was supplemented with ampicillin at 100 μg/mL for *E. coli*, carbenicillin at 250 μg/mL for *P. aeruginosa*, or tetracycline at 60 μg/mL for *E. coli* and 100 μg/mL for *P. aeruginosa* or gentamicin at 200 μg/mL for *P. aeruginosa* (Table S2). When necessary, UQ_4_ or 0.1% (w/v) arabinose final concentration was added to the medium to enhance bacterial anaerobic growth or to induce *P*_BAD_ expression of pSW196-derived plasmids, respectively. *Pseudomonas* Isolation Agar (PIA) medium (from DB) containing irgasan (25 μg/L) was used for triparental mating to counter select *E. coli*. For CFU counting, bacteria were suspended in PBS and cell suspensions were serially diluted in PBS. For each sample, 100 μL of at least four different dilutions were plated on LB plates, incubated for 24 h at 37°C and CFU were counted using a Scan 100 Interscience.

### Plasmids and genetic manipulations

The plasmids and the primers used in this study are listed in Tables S2 and S4). All the plasmids produced in this work were checked using DNA sequencing (GATC Biotech, Konstanz, Germany). To generate *P. aeruginosa* deletion mutants, overlapping upstream and downstream flanking regions of *ubiT_Pa_, ubiU_Pa_* and *ubiV_Pa_* genes were obtained by PCR amplification using PAO1 genome as template and the oligonucleotides described in Table S4. The resulting fragments were then cloned into *Sma*I-cut pEXG2 plasmid by sequence and ligation-independent cloning (45). To complement the mutants, the *ubiT_Pa_, ubiU_Pa_* and *ubiV_Pa_* fragments were generated by PCR amplification using the oligonucleotide pairs ubiT-PA-F/ubiT-PA-R, ubiU-PA-F/ubiU-PA-R and ubiV-PA-F/ubiV-PA-R, respectively, and PAO1 genome as template (Table S4). The fragments were *Eco*RI-*Sac*I digested and inserted into *P*_BAD_-harboring pSW196 plasmid, yielding the p*ubiT_Pa_*, p*ubiU_Pa_* and p*ubiV_Pa_* plasmids, respectively (Table S2). The pEXG2- and pSW196-derived vectors were transferred into *P. aeruginosa* PAO1 strain by triparental mating using pRK2013 as a helper plasmid (46). For allelic exchange using the pEXG2 plasmids, co-integration events were selected on PIA (*Pseudomonas* Isolation Agar) plates containing gentamicin. Single colonies were then cultured on NaCl-free LB agar plates containing 10% (w/vol) sucrose to select for the loss of the plasmid, and the resulting sucrose-resistant colonies were checked for mutant genotype by PCR. To overproduce C-terminally His-tagged UbiV_Pa_, the *ubiV_Pa_* gene was cloned into the pET22b(+) vector. The *ubiV_Pa_* insert was obtained by PCR amplification using the oligonucleotide pair pET22-UbiV-F and pET22-UbiV-R and *ubiV_Pa_* ORF as template (Table S4). *Nde*I-*Xho*I digested amplicon was ligated to *Nde*I-*Xho*I digested pET22b(+) vector to obtain pET22-UbiV_Pa_ (Table S2). Variants of UbiV_Pa_ were obtained using the Q5 Site-Directed Mutagenesis Kit (New England Biolabs) (UbiV_Pa_ C193AC197A and UbiV_Pa_ C39AC193AC197A) and the QuickChange II XL Site-Directed Mutagenesis Kit (Agilent) (UbiV_Pa_ C39AC180A) according to the manufacturer’s specifications using pET22b-UbiV_Pa_ as template (Table S2 and S4). The *ubiU_Pa_* gene was cloned into pET-22b(+) by following the same protocol as that for *ubiV_Pa_* gene. The *ubiT_Pa_* gene was synthesized by Eurofins with *E. coli* codon optimization. The synthetic gene was then cloned into the EcoRI/NotI sites of vector pET32a(+) (Novagen), resulting in plasmid pET32-TrxA-UbiT_Pa_ (Table S2).

### Soft-agar study to evaluate the O_2_-dependency of growth

Soft-agar studies were performed in denitrification medium supplemented by agar 0.7% (w/v) final concentration. After argon bubbling (O_2_<0.1 ppm) for 30 min, the suspension (13 mL) was autoclaved in Hungate tubes. They were then placed in a 40°C incubator and they were inoculated through the septum with 100 μL of overnight cultures taken with disposable syringe and needles from Eppendorf tubes filled to the top, mixed by inverting, and incubated at room temperature 30 min to allow the agar to solidify. Then, the tubes were incubated under aerobic conditions with caps loosened at 37°C for 24 h. A control experiment was performed with resazurin (0.25 μg/mL final concentration) used as an indicator of medium oxygenation. When required, the medium was supplemented with antibiotics.

### Lipid extractions and quinone analysis

Cultures (5 mL under ambient air and 13 mL under anaerobic conditions) were cooled down on ice before centrifugation at 3200 g, 4°C, 10 min. Cell pellets were washed in 1 mL ice-cold PBS and transferred to pre-weighted 1.5 mL Eppendorf tubes. After centrifugation at 12 000 g, 4°C, 1 min and elimination of supernatant, the cell wet weight was determined (~5-30 mg) and pellets were stored at −20°C. Quinone extraction from cell pellets was performed as previously described (18). Lipid extracts corresponding to 1 mg of cell wet weight were analyzed by HPLC electrochemical detection-mass spectrometry (ECD-MS) with a BetaBasic-18 column at a flow rate of 1 mL/min with mobile phases composed of methanol, ethanol and a mix of 90% isopropanol, 10% ammonium acetate (1 M), 0.1% TFA: mobile phase 1 (50% methanol, 40% ethanol and 10% mix). When necessary, MS detection was performed on a MSQ spectrometer (Thermo Scientific) with electrospray ionization in positive mode (probe temperature 400°C, cone voltage 80V). Single ion monitoring (SIM) detected the following compounds: UQ_8_ (M_+_NH_4_^+^), m/z 744-745, 6-10 min, scan time 0.2 s; UQ_9_ (M_+_NH_4_^+^), m/z 812-813, 9-14 min, scan time 0.2 s; UQ_10_ (M_+_NH_4_^+^), m/z 880.2-881.2, 10-17 min, scan time 0.2 s., DMQ_8_ (M_+_NH_4_^+^), m/z 714-715, 5-10 min, scan time 0.2 s. MS spectra were recorded between m/z 600 and 900 with a scan time of 0.3 s. ECD and MS peak areas were corrected for sample loss during extraction on the basis of the recovery of the UQ_10_ internal standard and were then normalized to cell wet weight. The peaks of UQ_8_ and UQ_9_ obtained with electrochemical detection were quantified with a standard curve of UQ_10_ as previously described (18).

### Overproduction and purification of UbiV, UbiU and UbiT from P. aeruginosa in E. coli

Wild-type and variants UbiV_Pa_ were expressed and purified as previously described for *E. coli* proteins (9). Briefly, the pET-22b(+) plasmid, encoding wild-type or variants UbiV_Pa_, were co-transformed with pGro7 plasmid (Takara Bio Inc.) into *E. coli* BL21 (DE3) Δ*ubiUV* competent cells grown at 37 °C in LB medium, which was supplemented with ampicillin (50 μg/mL), kanamycin (50 μg/mL) and chloramphenicol (12.5 μg/mL). At an OD_600_ = 1.2, D-arabinose was added to the cultures at a final concentration of 2 mg/mL. At an OD_600_ = 1.8, cultures were cooled down on ice for 20 min, and IPTG was added at a final concentration of 0.1 mM. Cells were then allowed to grow further at 16 °C overnight. Wild-type UbiV_Pa_ and the different variants were purified by Ni-NTA chromatography followed by SEC in buffer A (50 mM Tris-HCl, 25 mM NaCl, 15% (v/v) glycerol, pH 8.5) containing 1 mM DTT. The purified proteins were concentrated to 30-40 mg/mL using Amicon concentrators (30-kDa cutoff; Millipore).

The overproduction of wild-type UbiU_Pa_ was performed in *E. coli* BL21 (DE3) Δ*ubiUV* cells by following the same protocol as that for UbiV_Pa_, except that UbiU_Pa_ over-expression was induced with 0.05 mM of IPTG and the cell pellets were resuspended in buffer B (50 mM Tris-HCl, 500 mM NaCl, 15% (v/v) glycerol, pH 8.5) containing 0.2% (w/v) N-lauroylsarcosine sodium salt. After cell disruption by sonication, the clarified cell-free extracts were loaded onto a His-Trap FF crude column (GE Healthcare) preequilibrated with buffer B containing 0.1% (w/v) N-lauroylsarcosine sodium salt. The column was washed with 10 column volumes of buffer C (50 mM Tris-HCl, 500 mM NaCl, 15% (v/v) glycerol, 10 mM imidazole, pH 8.5) containing 6 mM N,N-dimethyldodecylamine N-oxide (LDAO) to remove non-specifically bound *E. coli* proteins and then eluted with a linear gradient of 10 column volumes of buffer C containing 500 mM imidazole and 6 mM LDAO. Fractions containing UbiU_Pa_ were pooled and then loaded on a HiLoad 16/600 Superdex 200 pg (GE Healthcare) pre-equilibrated in buffer D (50 mM Tris-HCl, 150 mM NaCl, 15% (v/v) glycerol, pH 8.5) containing 3 mM LDAO. The purified proteins were concentrated using Amicon concentrators (100-kDa cutoff; Millipore), aliquoted, frozen in liquid nitrogen, and stored at −80°C. For protein-lipid overlay, fractions 34-43 were pooled.

Overproduction of UbiT_Pa_ fused with the thioredoxin (TrxA-UbiT_Pa_) in *E. coli* BL21 (DE3) Δ*ubiUV* cells was performed by following the same protocol as that for UbiU_Pa_, except that over-expression of chimeric gene was induced at an OD_600_ of 0.5 and the cell pellet was resuspended in buffer B containing 5% (w/v) sodium cholate. TrxA-UbiT_Pa_ was first purified, following the same protocol as that for UbiU_Pa_, by Ni-NTA chromatography, except that the HisTrap FF crude column was pre-equilibrated with buffer B containing 0.5% (w/v) sodium cholate and then eluted with buffer C containing 500 mM imidazole and 0.5% (w/v) sodium cholate. Fractions containing TrxA-UbiT_Pa_ were pooled and detergent was removed using a Hiprep 26/10 Desalting column (GE Healthcare) preequilibrated with buffer D. The fusion protein was digested with thrombin (10 units/mg of TrxA-UbiT_Pa_) at room temperature and then loaded on a HiLoad 16/600 Superdex 200 pg (GE Healthcare) coupled with a HisTrap FF crude column (GE Healthcare) pre-equilibrated with buffer D. The purified proteins were concentrated using Amicon concentrators (100-kDa cutoff; Millipore), aliquoted, frozen in liquid nitrogen, and stored at −80°C.

### [Fe-S] cluster reconstitution

The [Fe-S] cluster(s) of holo-UbiV_Pa_ and holo-variants was reconstituted as previously described (9). Briefly, a solution containing 100 μM of as-purified proteins was treated with 5 mM DTT for 15 min at 20°C and then incubated for 1 h with a 5-fold molar excess of both ferrous ammonium sulfate and L-cysteine. The reaction was initiated by the addition of a catalytic amount of the *E. coli* cysteine desulfurase CsdA (1-2% molar equivalent) and monitored by UV-visible absorption spectroscopy. After 1 h of incubation, the holo-proteins were then loaded onto a Superdex 75 Increase 10/300 GL column (GE Healthcare) pre-equilibrated with buffer A. The fractions containing the holo-proteins were pooled and concentrated to 20-30 mg/mL on a Vivaspin concentrator (30-kDa cutoff).

### Protein–lipid overlay

To assess the lipid-binding properties of UbiT_Pa_ and UbiU_Pa_, a protein–lipid overlay was performed as previously described (47). Briefly, 2 μL of 20 mM lipids in dichloromethane were spotted onto PVDF membrane and allowed to dry at room temperature for 1 h. The membranes were blocked in 3 % (w/v) fatty acid-free BSA in TBST (50 mM Tris-HCl, 150 mM NaCl and 0.1% (v/v) Tween-20, pH 7.5) for 1 h. The membranes were then incubated overnight at 4 °C with gentle stirring in the same solution containing 0.2 μg/mL of the indicated proteins. After washing six times during 30 min in TBST buffer, the membranes were incubated for 1 h with a 1/1000 dilution of anti-polyHis monoclonal antibody (Sigma) and then for 1 h with a 1/10 000 dilution of anti-mouse– horseradish peroxidase conjugate (Thermo Fisher Scientific). His-tagged proteins bound to the membrane by virtue of its interaction with lipid, were detected by enhanced chemiluminescence using Clarity Max Western ECL Substrate (Bio-rad).

### Quantification methods

Protein concentrations were determined using the method of Bradford (Bio-Rad) with bovine serum albumin as the standard. The iron and acid-labile sulfide were determined according to the method of Fish (48) and Beinert (49), respectively, before and after [4Fe-4S] cluster reconstitution.

### UV-vis spectroscopy

UV-visible spectra were recorded in 1-cm optic path quartz cuvettes under aerobic conditions on a Cary 100 Uv-vis spectrophotometer (Agilent) and under anaerobic conditions in a glove box on a XL-100 Uvikon spectrophotometer equipped with optical fibers.

## The abbreviations used are

MK, menaquinone; UQ, ubiquinone; DMK, dimethyl-menaquinone; DMQ, 2-octaprenyl-3-methyl-6-methoxy-1,4-benzoquinone; SEC, size exclusion chromatography; SCP2, sterol carrier protein 2; IPTG, isopropyl-1-thio-β-D-galactopyranoside; ECD, electrochemical detection; PA: phosphatidic acid; POPE: 3-methylcatechol, 1-palmitoyl-2-oleoyl-*sn*-glycero-3-phosphoethanolamine.

## Author contributions

L. P., F. B., M. F., M. L. conceived and designed the experiments. C. -D. -T., J. M., S. E., E. B., and L. P. performed the experiments. E. B. and B. F. contributed reagents/materials/analysis tools. L. P., M. L., and F. P. wrote the paper. All authors analyzed the results and approved the final version of the manuscript.

## Acknowledgements

This work was supported by the Agence Nationale de la Recherche (ANR), projects (An)aeroUbi ANR-15-CE11-0001-02, O_2_-taboo ANR-19-CE44-0014, DYNAMO ANR-11-LABX-0011-01 and ANR-10-LABX-62-IBEID, the University Grenoble Alpes (UGA), the French Centre National de la Recherche Scientifique (CNRS) and the Commissariat à l’Energie Atomique et aux Energies Alternatives (CEA)

## Data availability

All data is contained within the manuscript and supplemental materials.

---

This article contains Tables S1 to S4 and Figures S1 to S6.

**Table S1:** Bioinformatics analysis of the genes required for UQ synthesis in *E. coli* and *P. aeruginosa*.

| *E. coli* MG1655 |  |  |  | *P. aeruginosa* PA01 |  |  | Functions in *E. coli* (From https://ecocyc.org/.) or specific to *P. aeruginosa* |
| --- | --- | --- | --- | --- | --- | --- | --- |
| Gene ID | Size (aa) | Need in UQ biosynthesis (1) |  | Gene ID | Size (aa) | % of identity |  |
|  |  | +O<sub>2</sub> | -O<sub>2</sub> |  |  |  |  |
| *ispB* | 323 | Yes |  | *PA4569* | 321 | 52 | *all-trans*-octaprenyl-diphosphate synthase |
| *ubiA* | 290 | Yes |  | *PA5358* | 296 | 56 | 4-HB octaprenyl transferase |
| *ubiB* | 546 | Yes |  | *PA5065* | 533 | 56 | Essential to UQ biosynthesis |
| *ubiC* | 165 | Yes |  | *PA5357* | 178 | 31 | Chorismate pyruvate lyase |
| *ubiD* | 497 | Yes |  | *PA5237* | 496 | 77 | 3-octaprenyl 4-HB decarboxylase |
| *ubiE* | 251 | Yes |  | *PA5063* | 256 | 73 | 2-octaprenyl-6-methoxy-1,4-benzoquinone methylase |
| *ubiF* | 391 | Yes |  | No homologue |  |  | 2-octaprenyl-3-methyl-6-methoxy-1,4-benzoquinol hydroxylase |
| *coq7* | No homologue |  |  | *PA0655* | 116 | - | 2-nonaprenyl-3-methyl-6-methoxy-1,4-benzoquinol hydroxylase (2) |
| *ubiG* | 240 | Yes |  | *PA3171* | 232 | 64 | 3-demethylubiquinone-8 3-*O*-methyl transferase<br>2-octaprenyl-6-hydroxyphenol methylase |
| *ubiH* | 392 | Yes | No | *PA5223* | 394 | 40 | 2-octaprenyl-6-methoxyphenol hydroxylase |
| *ubiI* | 400 | Yes | No | *PA5221* | 405 | 45 | 2-octaprenylphenol hydroxylase |
| *ubiJ* | 201 | Yes | No | *PA5064* | 208 | 21 | Essential to UQ biosynthesis, SCP2 domain |
| *ubiK* | 96 | Yes | No | *PA5289* | 86 | 37 | Ubiquinone biosynthesis accessory factor |
| *ubiT* | 174 | No | Yes | *PA3911* | 171 | 42 | Essential to UQ biosynthesis, SCP2 domain |
| *ubiU* | 331 | No | Yes | *PA3913* | 331 | 64 | Essential to UQ |
| *ubiV* | 292 | No | Yes | *PA3912* | 296 | 50 | Essential to UQ |
| *ubiX* | 189 | Yes |  | *PA4019* | 209 | 40 | Flavin prenyltransferase - UbiD partner protein |

**Table S2:**
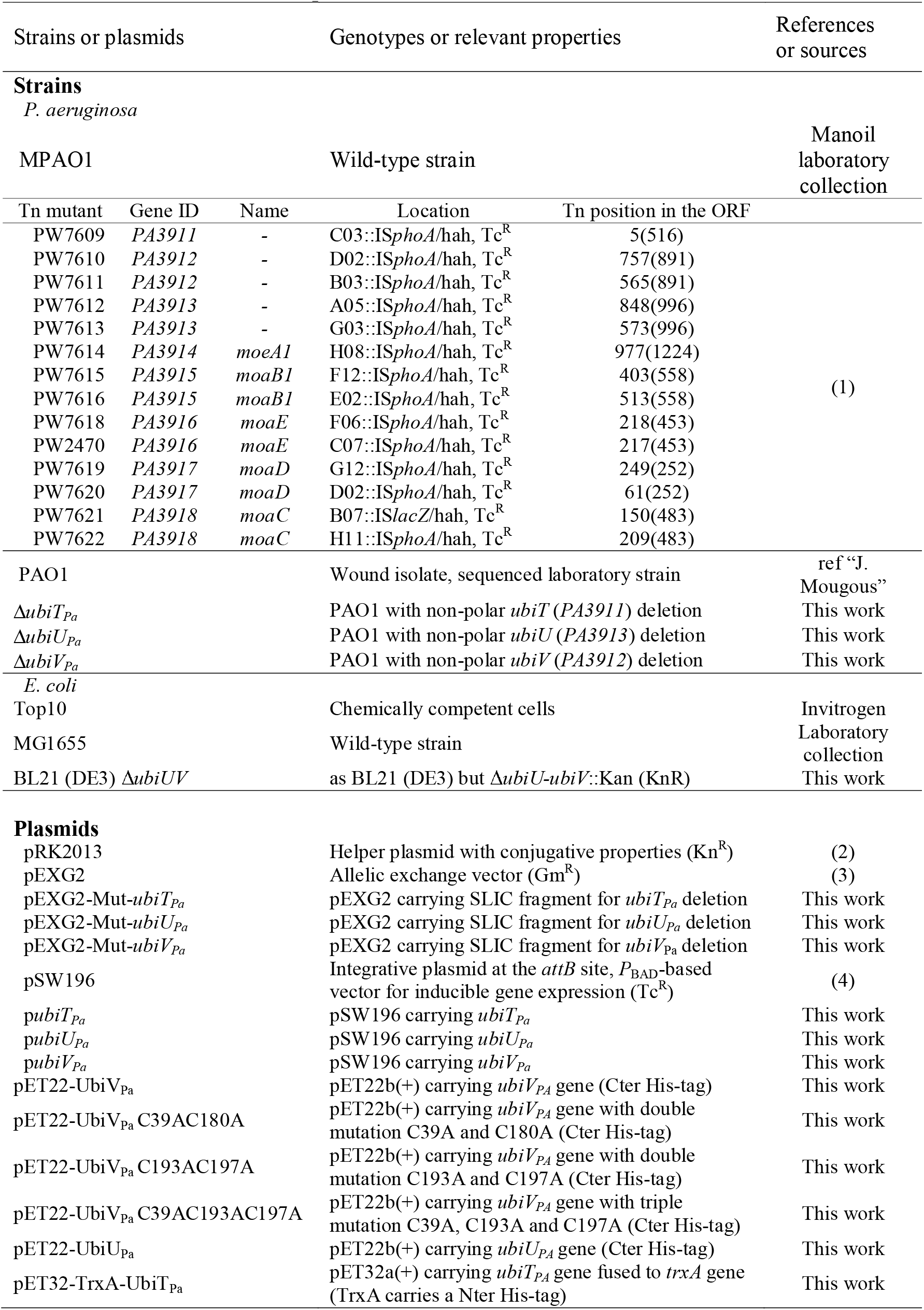
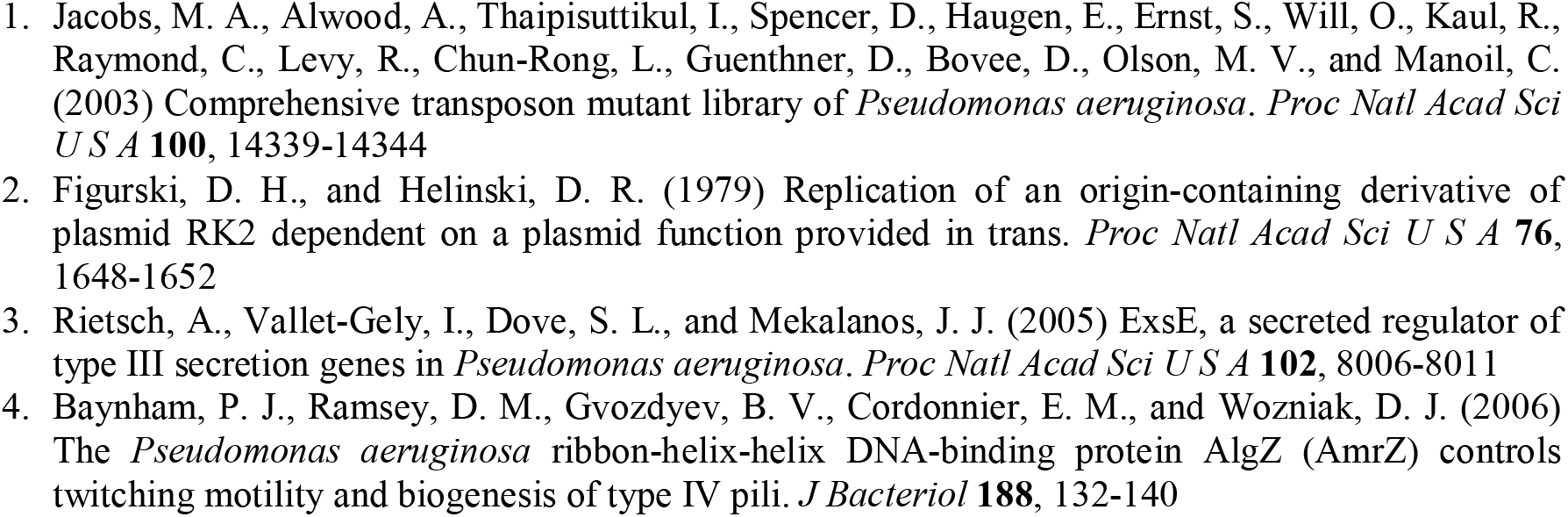
Bacterial strains and plasmids used in this work.

**Table S3:** isoprenoid quinones content of the SEC-purified UbiPa proteins.

| Purified proteins | UQ<sub>8</sub> (pmol/mg proteins) | DMQ<sub>8</sub> (AU/mg proteins) |
| --- | --- | --- |
| UbiV<sub>Pa</sub> | nd | nd |
| UbiT<sub>Pa</sub><sup>1</sup> | 9.75±4.57 | nd |
| UbiU<sub>Pa</sub> (39)<sup>2</sup> | 488 | 19932 |
| UbiU<sub>Pa</sub> (40)<sup>2</sup> | 457 | 20603 |

**Table S4:**
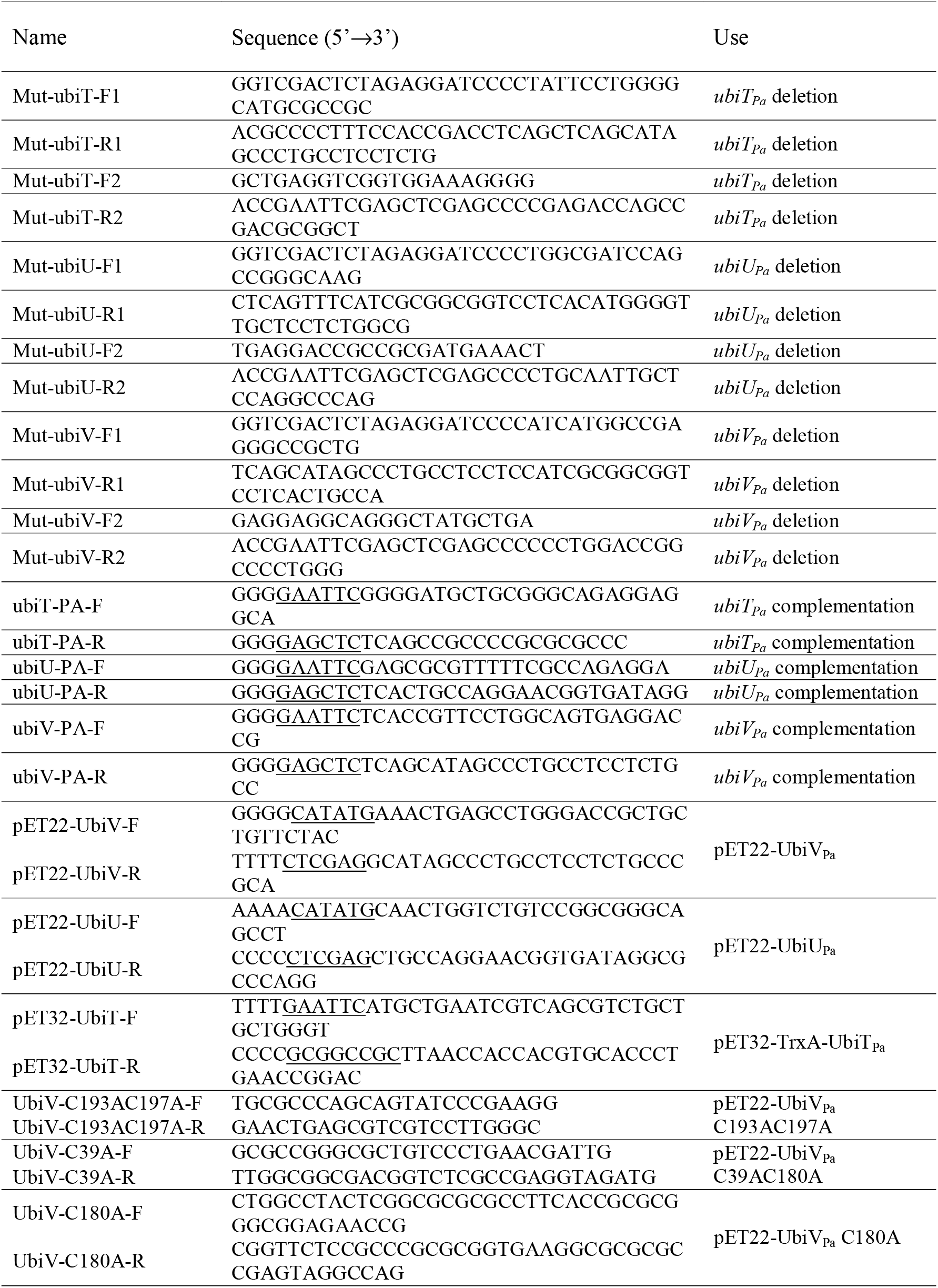
Primers used in this work. Restriction sites are underlined.

**Figure S1:**
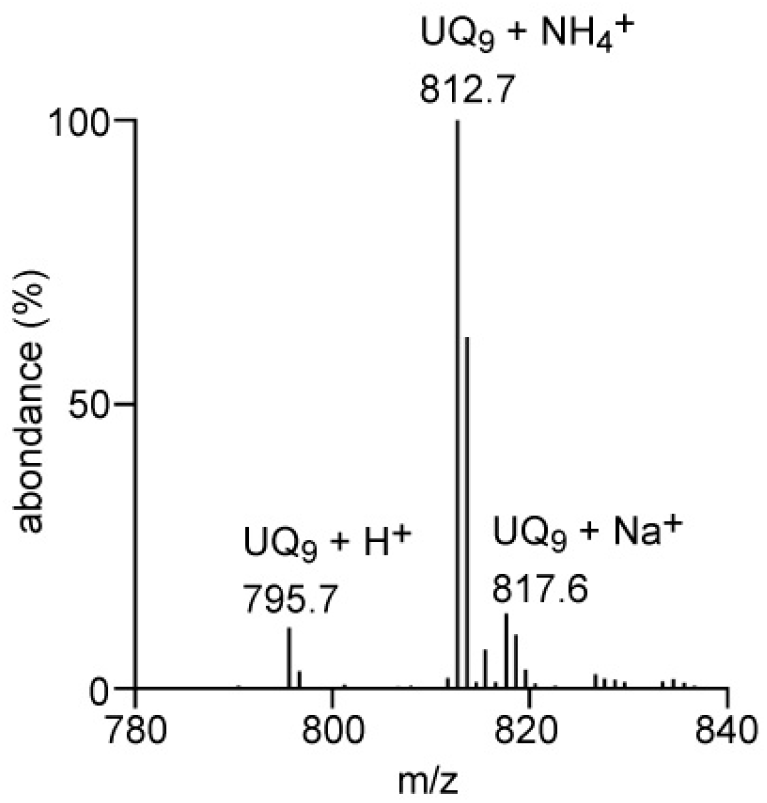
Zoom (740 to 840 m/z region) of the mass spectrum displayed in Fig. 1*C*. H^+^, NH4^+^ and Na^+^ adducts corresponding to UQ_9_ were indicated.

**Figure S2:**
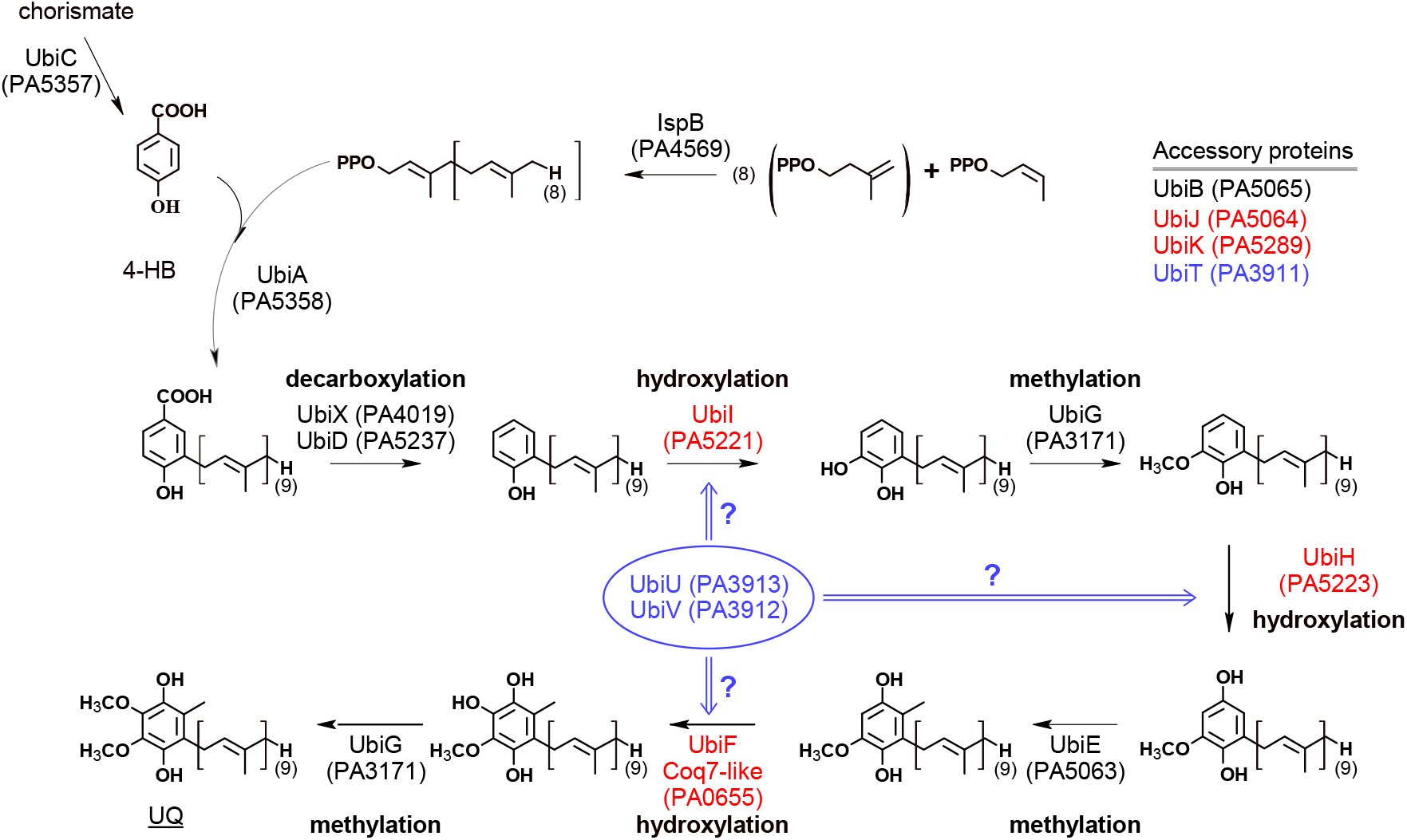
Putative UQ_9_ biosynthetic pathways in *E. coli* MG1655 and in *P. aeruginosa* PAO1. Proteins common to both biosynthetic pathways (aerobic and anaerobic) are indicated in black. Proteins involved only in aerobiosis or in anaerobiosis are indicated in red or in blue, respectively. The function of UbiT, UbiU and UbiV remains unclear. Corresponding gene IDs in *P. aeruginosa* are indicated in parenthesis. UbiF is only identified in *E. coli* and its functional homolog in *P. aeruginosa* is a Coq7-like hydroxylase.

**Figure S3:**
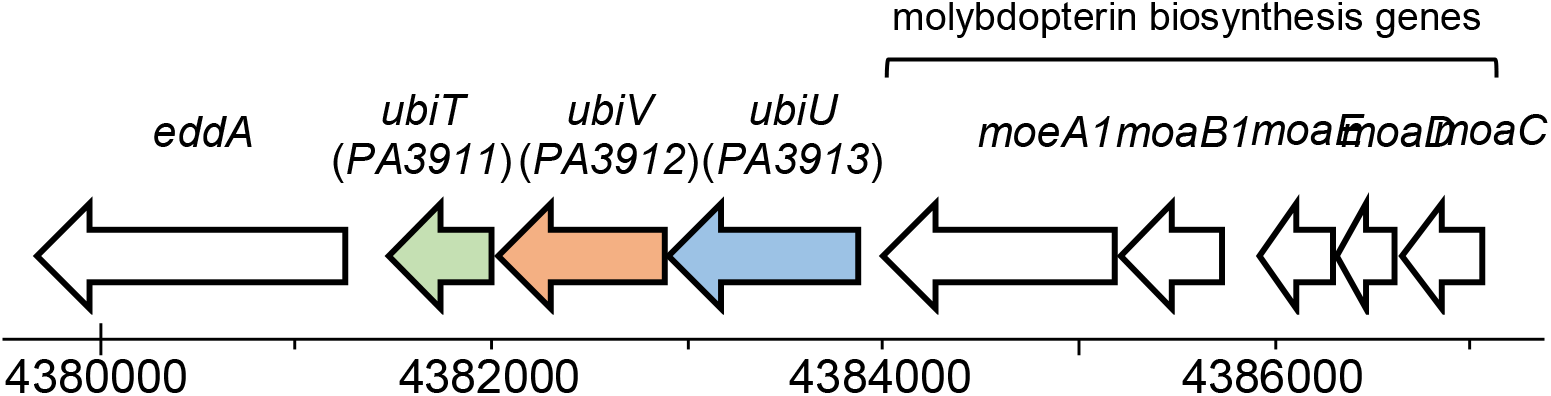
Genomic localization of the *ubiUVT* operon in *P. aeruginosa* PAO1. ORFs of the genes *ubiT_Pa_, ubiU_Pa_* and *ubiV_Pa_* are represented by green, blue and orange arrows, respectively.

**Figure S4:**
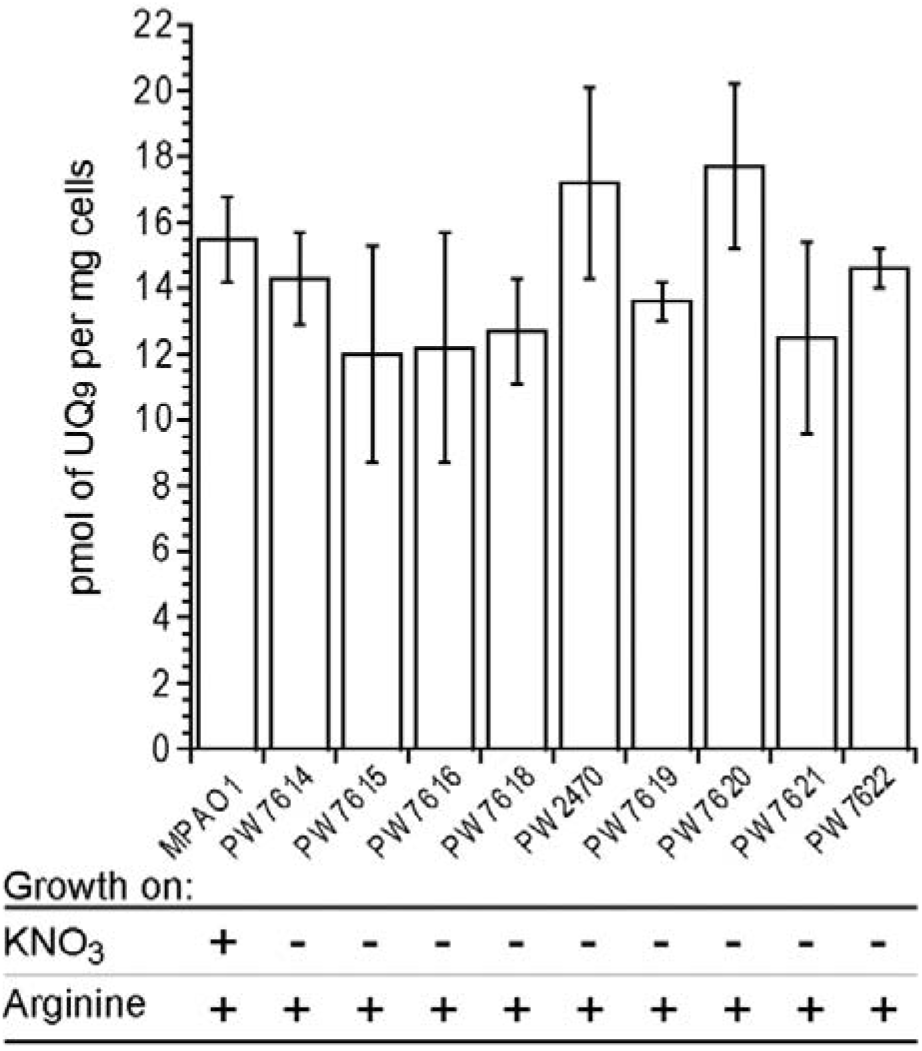
Molybdopterin cofactors are not involved in anaerobic biosynthesis of UQ_9_ in *P. aeruginosa.* Quantification of cellular UQ_9_ content (*n=*3) in lipid extracts from wild-type MPAO1 and Tn mutants cells (see Table S2) grown 48h anaerobically in LB medium supplemented with arginine (40 mM final concentration). Error bars are means ± S.D. Anaerobic growth of wild-type and Tn mutants was assessed in denitrification medium or in LB supplemented with arginine (40 mM final concentration) after 48h of incubation. (+), growth; (-) no growth.

**Figure S5:**
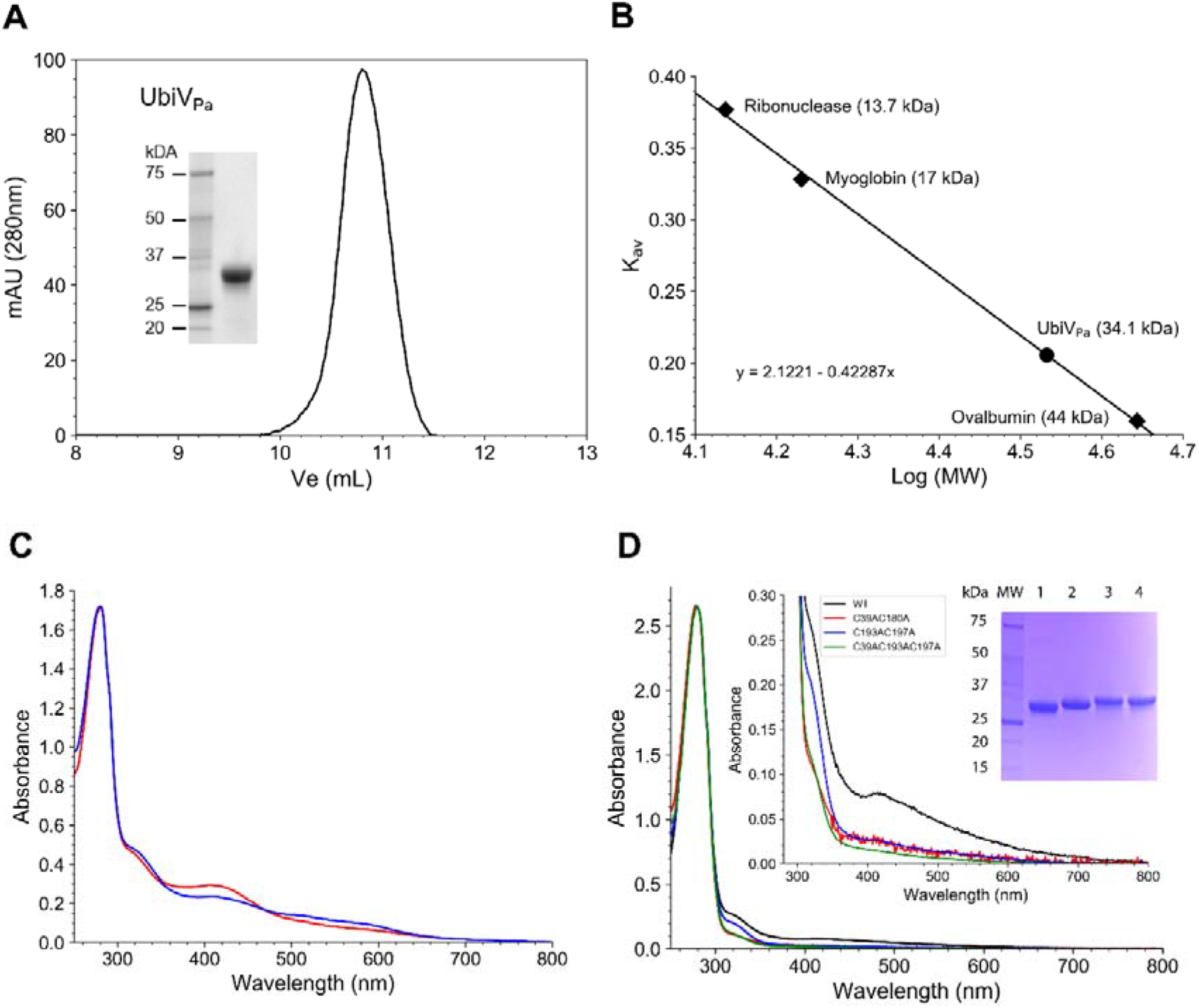
Spectral characterization of recombinant ubiV_Pa_ and its variants. *A*, Gel filtration chromatogram of aerobically purified ubiV_Pa_ on a Superdex 75 Increase 10/300 GL column; Inset: SDS-PAGE of purified ubiV_Pa_. *B*, Calibration curve, standard proteins in rhombi, ubiV_Pa_ in circle. *C*, UV-visible absorption spectra of 22.7 μM holo-UbiV_Pa_ under anaerobiosis (red) and after 30 min exposure to air (blue). *D*, Comparative UV-visible absorption spectra of aerobically purified wild-type (black) and various Cys-to-Ala mutants of ubiV_Pa_ (C39AC180A in red; C193AC197A in blue; C39AC193AC197A in green). Inset: enlargement of the 300 to 800 nm region and Coomassie staining SDS-PAGE showing the purifications of WT-UbiV_Pa_ and its variants. Lanes: Mw, molecular weight markers; 1, wild-type ubiV_Pa_; 2, ubiV_Pa_ C39AC180A; 3, ubiV_Pa_ C193AC197A; 4, ubiV_Pa_ C39AC193AC197A. All proteins were suspended in 50 mM Tris-HCl, 25 mM NaCl, 15% (v/v) glycerol, 1 mM DTT, pH 8.5.

**Figure S6:**
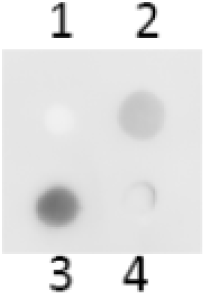
Protein-lipid overlay assay between recombinant ubiT_Pa_ and phosphatidic acid (PA). 2 μL of four different lipid/compound (1, 3-methylcatechol; 2, POPE; 3, UQ_10_ and 4, cholesterol) at 20 mM final concentration were spotted on a PVDF membrane and then incubated with ubiT_Pa_ at 0.2 μg/mL final concentration.

## REFERENCES

1. Crull, M. R., Somayaji, R., Ramos, K. J., Caldwell, E., Mayer-Hamblett, N., Aitken, M. L., Nichols, D. P., Rowhani-Rahbar, A., and Goss, C. H. (2018) Changing Rates of Chronic *Pseudomonas aeruginosa* Infections in Cystic Fibrosis: A Population-Based Cohort Study. Clin Infect Dis 67, 1089–1095

2. Magill, S. S., Edwards, J. R., Bamberg, W., Beldavs, Z. G., Dumyati, G., Kainer, M. A., Lynfield, R., Maloney, M., McAllister-Hollod, L., Nadle, J., Ray, S. M., Thompson, D. L., Wilson, L. E., Fridkin, S. K., Emerging Infections Program Healthcare-Associated, I., and Antimicrobial Use Prevalence Survey, T. (2014) Multistate point-prevalence survey of health care-associated infections. N Engl J Med 370, 1198–1208

3. Arai, H. (2011) Regulation and Function of Versatile Aerobic and Anaerobic Respiratory Metabolism in *Pseudomonas aeruginosa*. Front Microbiol 2, 103

4. Williams, H. D., Zlosnik, J. E., and Ryall, B. (2007) Oxygen, cyanide and energy generation in the cystic fibrosis pathogen *Pseudomonas aeruginosa*. Adv Microb Physiol 52, 1–71

5. Torres, A., Kasturiarachi, N., DuPont, M., Cooper, V. S., Bomberger, J., and Zemke, A. (2019) NADH Dehydrogenases in *Pseudomonas aeruginosa* Growth and Virulence. Front Microbiol 10, 75

6. Page, A. C., Jr., Gale, P., Wallick, H., Walton, R. B., Mc, D. L., Woodruff, H. B., and Folkers, K. (1960) Coenzyme Q. 17. Isolation of coenzyme Q10 from bacterial fermentation. Arch Biochem Biophys 89, 318–321

7. Matsushita, K., Yamada, M., Shinagawa, E., Adachi, O., and Ameyama, M. (1980) Function of ubiquinone in the electron transport system of *Pseudomonas aeruginosa* grown aerobically. J Biochem 88, 757–764

8. Nowicka, B., and Kruk, J. (2010) Occurrence, biosynthesis and function of isoprenoid quinones. Biochim Biophys Acta 1797, 1587–1605

9. Pelosi, L., Vo, C. D., Abby, S. S., Loiseau, L., Rascalou, B., Hajj Chehade, M., Faivre, B., Gousse, M., Chenal, C., Touati, N., Binet, L., Cornu, D., Fyfe, C. D., Fontecave, M., Barras, F., Lombard, M., and Pierrel, F. (2019) Ubiquinone Biosynthesis over the Entire O2 Range: Characterization of a Conserved O2-Independent Pathway. mBio 10

10. Alexander, K., and Young, I. G. (1978) Alternative hydroxylases for the aerobic and anaerobic biosynthesis of ubiquinone in *Escherichia coli*. Biochemistry 17, 4750–4755

11. Hajj Chehade, M., Pelosi, L., Fyfe, C. D., Loiseau, L., Rascalou, B., Brugiere, S., Kazemzadeh, K., Vo, C. D., Ciccone, L., Aussel, L., Coute, Y., Fontecave, M., Barras, F., Lombard, M., and Pierrel, F. (2019) A Soluble Metabolon Synthesizes the Isoprenoid Lipid Ubiquinone. Cell Chem Biol 26, 482–492 e487

12. Zumft, W. G. (1997) Cell biology and molecular basis of denitrification. Microbiol Mol Biol Rev 61, 533–616

13. Jensen, P. O., Kolpen, M., Kragh, K. N., and Kuhl, M. (2017) Microenvironmental characteristics and physiology of biofilms in chronic infections of CF patients are strongly affected by the host immune response. APMIS 125, 276–288

14. Borrero-de Acuna, J. M., Timmis, K. N., Jahn, M., and Jahn, D. (2017) Protein complex formation during denitrification by *Pseudomonas aeruginosa*. Microb Biotechnol 10, 1523–1534

15. Stenmark, P., Grunler, J., Mattsson, J., Sindelar, P. J., Nordlund, P., and Berthold, D. A. (2001) A new member of the family of di-iron carboxylate proteins. Coq7 (clk-1), a membrane-bound hydroxylase involved in ubiquinone biosynthesis. J Biol Chem 276, 33297–33300

16. Jiang, H. X., Wang, J., Zhou, L., Jin, Z. J., Cao, X. Q., Liu, H., Chen, H. F., and He, Y. W. (2019) Coenzyme Q biosynthesis in the biopesticide Shenqinmycin-producing *Pseudomonas aeruginosa* strain M18. J Ind Microbiol Biotechnol

17. Coelho, C., and Romao, M. J. (2015) Structural and mechanistic insights on nitrate reductases. Protein Sci 24, 1901–1911

18. Hajj Chehade, M., Loiseau, L., Lombard, M., Pecqueur, L., Ismail, A., Smadja, M., Golinelli-Pimpaneau, B., Mellot-Draznieks, C., Hamelin, O., Aussel, L., Kieffer-Jaquinod, S., Labessan, N., Barras, F., Fontecave, M., and Pierrel, F. (2013) ubiI, a new gene in *Escherichia coli* coenzyme Q biosynthesis, is involved in aerobic C5-hydroxylation. J Biol Chem 288, 20085–20092

19. Loiseau, L., Fyfe, C., Aussel, L., Hajj Chehade, M., Hernandez, S. B., Faivre, B., Hamdane, D., Mellot-Draznieks, C., Rascalou, B., Pelosi, L., Velours, C., Cornu, D., Lombard, M., Casadesus, J., Pierrel, F., Fontecave, M., and Barras, F. (2017) The UbiK protein is an accessory factor necessary for bacterial ubiquinone (UQ) biosynthesis and forms a complex with the UQ biogenesis factor UbiJ. J Biol Chem 292, 11937–11950

20. Hille, R. (2005) Molybdenum-containing hydroxylases. Arch Biochem Biophys 433, 107–116

21. Filiatrault, M. J., Picardo, K. F., Ngai, H., Passador, L., and Iglewski, B. H. (2006) Identification of *Pseudomonas aeruginosa* genes involved in virulence and anaerobic growth. Infect Immun 74, 4237–4245

22. Imlay, J. A. (2006) Iron-sulphur clusters and the problem with oxygen. Mol Microbiol 59, 1073–1082

23. Ollagnier de Choudens, S., and Barras, F. (2017) Genetic, Biochemical, and Biophysical Methods for Studying Fe-S Proteins and Their Assembly. Methods Enzymol 595, 1–32

24. Ollagnier-de Choudens, S., Loiseau, L., Sanakis, Y., Barras, F., and Fontecave, M. (2005) Quinolinate synthetase, an iron-sulfur enzyme in NAD biosynthesis. FEBS Lett 579, 3737–3743

25. Groenewold, M. K., Massmig, M., Hebecker, S., Danne, L., Magnowska, Z., Nimtz, M., Narberhaus, F., Jahn, D., Heinz, D. W., Jansch, L., and Moser, J. (2018) A phosphatidic acid-binding protein is important for lipid homeostasis and adaptation to anaerobic biofilm conditions in *Pseudomonas aeruginosa*. Biochem J 475, 1885–1907

26. Dowhan, W. (1997) Molecular basis for membrane phospholipid diversity: why are there so many lipids? Annu Rev Biochem 66, 199–232

27. Aussel, L., Pierrel, F., Loiseau, L., Lombard, M., Fontecave, M., and Barras, F. (2014) Biosynthesis and physiology of coenzyme Q in bacteria. Biochim Biophys Acta 1837, 1004–1011

28. Sharma, P., Teixeira de Mattos, M. J., Hellingwerf, K. J., and Bekker, M. (2012) On the function of the various quinone species in Escherichia coli. FEBS J 279, 3364–3373

29. Nitzschke, A., and Bettenbrock, K. (2018) All three quinone species play distinct roles in ensuring optimal growth under aerobic and fermentative conditions in *E. coli* K12. PLoS One 13, e0194699

30. Filiatrault, M. J., Wagner, V. E., Bushnell, D., Haidaris, C. G., Iglewski, B. H., and Passador, L. (2005) Effect of anaerobiosis and nitrate on gene expression in *Pseudomonas aeruginosa*. Infect Immun 73, 3764–3772

31. Wu, M., Guina, T., Brittnacher, M., Nguyen, H., Eng, J., and Miller, S. I. (2005) The *Pseudomonas aeruginosa* proteome during anaerobic growth. J Bacteriol 187, 8185–8190

32. Worlitzsch, D., Tarran, R., Ulrich, M., Schwab, U., Cekici, A., Meyer, K. C., Birrer, P., Bellon, G., Berger, J., Weiss, T., Botzenhart, K., Yankaskas, J. R., Randell, S., Boucher, R. C., and Doring, G. (2002) Effects of reduced mucus oxygen concentration in airway *Pseudomonas* infections of cystic fibrosis patients. J Clin Invest 109, 317–325

33. Kolpen, M., Hansen, C. R., Bjarnsholt, T., Moser, C., Christensen, L. D., van Gennip, M., Ciofu, O., Mandsberg, L., Kharazmi, A., Doring, G., Givskov, M., Hoiby, N., and Jensen, P. O. (2010) Polymorphonuclear leucocytes consume oxygen in sputum from chronic *Pseudomonas aeruginosa* pneumonia in cystic fibrosis. Thorax 65, 57–62

34. Kolpen, M., Bjarnsholt, T., Moser, C., Hansen, C. R., Rickelt, L. F., Kuhl, M., Hempel, C., Pressler, T., Hoiby, N., and Jensen, P. O. (2014) Nitric oxide production by polymorphonuclear leucocytes in infected cystic fibrosis sputum consumes oxygen. Clin Exp Immunol 177, 310–319

35. Linnane, S. J., Keatings, V. M., Costello, C. M., Moynihan, J. B., O’Connor, C. M., Fitzgerald, M. X., and McLoughlin, P. (1998) Total sputum nitrate plus nitrite is raised during acute pulmonary infection in cystic fibrosis. Am J Respir Crit Care Med 158, 207–212

36. Skurnik, D., Roux, D., Aschard, H., Cattoir, V., Yoder-Himes, D., Lory, S., and Pier, G. B. (2013) A comprehensive analysis of *in vitro* and *in vivo* genetic fitness of *Pseudomonas aeruginosa* using high-throughput sequencing of transposon libraries. PLoS Pathog 9, e1003582

37. Navais, R., Mendez, J., Perez-Pascual, D., Cascales, D., and Guijarro, J. A. (2014) The yrpAB operon of Yersinia ruckeri encoding two putative U32 peptidases is involved in virulence and induced under microaerobic conditions. Virulence 5, 619–624

38. Sakai, Y., Kimura, S., and Suzuki, T. (2019) Dual pathways of tRNA hydroxylation ensure efficient translation by expanding decoding capability. Nat Commun 10, 2858

39. Lauhon, C. T. (2019) Identification and Characterization of Genes Required for 5-Hydroxyuridine Synthesis in *Bacillus subtilis* and *Escherichia coli* tRNA. J Bacteriol 201

40. Aussel, L., Loiseau, L., Hajj Chehade, M., Pocachard, B., Fontecave, M., Pierrel, F., and Barras, F. (2014) ubiJ, a new gene required for aerobic growth and proliferation in macrophage, is involved in coenzyme Q biosynthesis in *Escherichia coli* and *Salmonella enterica* serovar *Typhimurium*. J Bacteriol 196, 70–79

41. Held, K., Ramage, E., Jacobs, M., Gallagher, L., and Manoil, C. (2012) Sequence-verified two-allele transposon mutant library for *Pseudomonas aeruginosa* PAO1. J Bacteriol 194, 6387–6389

42. Jacobs, M. A., Alwood, A., Thaipisuttikul, I., Spencer, D., Haugen, E., Ernst, S., Will, O., Kaul, R., Raymond, C., Levy, R., Chun-Rong, L., Guenthner, D., Bovee, D., Olson, M. V., and Manoil, C. (2003) Comprehensive transposon mutant library of *Pseudomonas aeruginosa*. Proc Natl Acad Sci U S A 100, 14339–14344

43. Wagner, V. E., Bushnell, D., Passador, L., Brooks, A. I., and Iglewski, B. H. (2003) Microarray analysis of *Pseudomonas aeruginosa* quorum-sensing regulons: effects of growth phase and environment. J Bacteriol 185, 2080–2095

44. Hassett, D. J. (1996) Anaerobic production of alginate by *Pseudomonas aeruginosa:* alginate restricts diffusion of oxygen. J Bacteriol 178, 7322–7325

45. Jeong, J. Y., Yim, H. S., Ryu, J. Y., Lee, H. S., Lee, J. H., Seen, D. S., and Kang, S. G. (2012) One-step sequence-and ligation-independent cloning as a rapid and versatile cloning method for functional genomics studies. Appl Environ Microbiol 78, 5440–5443

46. Figurski, D. H., and Helinski, D. R. (1979) Replication of an origin-containing derivative of plasmid RK2 dependent on a plasmid function provided in trans. Proc Natl Acad Sci U S A 76, 1648–1652

47. Kavran, J. M., Klein, D. E., Lee, A., Falasca, M., Isakoff, S. J., Skolnik, E. Y., and Lemmon, M. A. (1998) Specificity and promiscuity in phosphoinositide binding by pleckstrin homology domains. J Biol Chem 273, 30497–30508

48. Fish, W. W. (1988) Rapid colorimetric micromethod for the quantitation of complexed iron in biological samples. Methods in enzymology 158, 357–364

49. Beinert, H. (1983) Semi-micro methods for analysis of labile sulfide and of labile sulfide plus sulfane sulfur in unusually stable iron-sulfur proteins. Analytical biochemistry 131, 373–378

